# Mapping rhodopsin trafficking in rod photoreceptors with quantitative super-resolution microscopy

**DOI:** 10.1101/2023.04.20.537413

**Authors:** Kristen N. Haggerty, Shannon C. Eshelman, Lauren A. Sexton, Emmanuel Frimpong, Leah M. Rogers, Melina A. Agosto, Michael A. Robichaux

## Abstract

Photoreceptor cells in the vertebrate retina have a highly compartmentalized morphology for efficient long-term phototransduction. Rhodopsin, the visual pigment in rod photoreceptors, is densely packaged into the rod outer segment sensory cilium and continuously renewed through essential synthesis and trafficking pathways housed in the rod inner segment. Despite the importance of this region for rod health and maintenance, the subcellular organization of rhodopsin and its trafficking regulators in the mammalian rod inner segment remain undefined. We used super-resolution fluorescence microscopy with optimized retinal immunolabeling techniques to perform a single molecule localization analysis of rhodopsin in the inner segments of mouse rods. We found that a significant fraction of rhodopsin molecules was localized at the plasma membrane in an even distribution along the entire length of the inner segment, where markers of transport vesicles also colocalized. Thus, our results collectively establish a model of rhodopsin trafficking through the inner segment plasma membrane as an essential subcellular pathway in mouse rod photoreceptors.

**SUMMARY:** Photoreceptor cells of the retina are maintained through a complex protein trafficking network. This study applies quantitative super-resolution microscopy to uncover localization details about the trafficking of the essential visual pigment rhodopsin in the inner segment region of rod photoreceptors.

## INTRODUCTION

In the vertebrate retina, vision is initiated by the phototransduction cascade in the outer segment (OS) sensory cilia of rod and cone photoreceptor cells. The light-absorbing G-protein coupled receptor (GPCR) proteins rhodopsin (Rho) in rods and the cone opsins in cones are densely packaged into flattened membrane discs that are stacked within the OS cilium. In the mouse retina, each rod OS contains ∼800 discs (Liang et al., 2004) with a Rho packaging density of ∼75,000 Rho molecules per disc (Skiba et al., 2023; Lyubarsky et al., 2004; Nickell et al., 2007). Rod OS discs are completely renewed in about 10 days through the process of disc shedding from the distal rod OS tips (Young, 1967; Kevany and Palczewski, 2010). As such, new discs are formed at the OS base, and these nascent discs must be filled with newly-synthesized protein, primarily Rho, which is synthesized in the rod photoreceptor inner segment (IS) and trafficked to the OS by way of the connecting cilium (CC) (Wensel et al., 2016; May-Simera et al., 2017). In mouse rods, the CC is a thin ciliary bridge that spans 1.1 µm between the IS and OS and is composed of an axoneme core of 9 microtubule doublets that extend into the OS (Gilliam et al., 2012). A coordinated homeostasis of OS disc shedding, nascent disc formation, IS protein delivery and CC trafficking must be maintained in photoreceptors for a lifetime of proper vision.

Mislocalization of Rho is a hallmark phenotype reported in many animal models of retinal disease, including models for retinitis pigmentosa (RP) and other retinal ciliopathies. RP is an inherited neurodegeneration affecting 1:4,000 in the USA (Hamel, 2006) that causes neuronal cell death in rod photoreceptors and subsequent loss of vision. In animal models, a wide range of Rho RP mutations have been demonstrated to cause Rho protein mislocalization to the rod IS region, as well as some degree of Rho mislocalization to the photoreceptor outer nuclear layer (ONL) and outer plexiform layer (OPL) (Sung et al., 1994; Chuang et al., 2004; Tam and Moritz, 2006; Santhanam et al., 2020; Robichaux et al., 2022). RP mutations to other rod-specific genes result in Rho mislocalization, including in *tulp1^-/-^* and *Pde6b^rd10^* mouse models of autosomal recessive RP (Hagstrom et al., 1999; Grossman et al., 2009), as well as in *RPGR* (retinitis pigmentosa GTPase regulator) mutant mouse and dog models of X-linked RP (Beltran et al., 2006; Megaw et al., 2017). Mouse models for syndromic retinal ciliopathies, such as Bardet-Biedl syndrome (BBS), Joubert syndrome (JBTS) and Meckel-Gruber syndrome (MKS), are caused by mutations in cilia-related genes that lead to rod degeneration in the retina caused by defective ciliary ultrastructure that is comorbid with Rho mislocalization (Abd-El-Barr et al., 2007; Hsu et al., 2017; Dilan et al., 2019; Guo et al., 2022). Similarly, Leber congenital amaurosis (LCA) mutations that disrupt CC-localized proteins SPATA7 and CEP290 also result in Rho mislocalization to the rod IS and ONL in mouse models (Eblimit et al., 2015; Potter et al., 2021). Collectively, rod dystrophies caused by any number of disruptions to normal rod homeostasis lead to Rho mislocalization, such that restorative therapeutic strategies to treat these retinal diseases must account for proper trafficking of Rho to preserve long-term rod stability. Therefore, a thorough understanding of the cellular mechanisms of Rho synthesis and trafficking in the IS of rods is essential for the development of effective retinal therapies.

The rod IS compartment is the biosynthetic domain of rods that is filled with endoplasmic reticulum (ER) and Golgi secretory organelles, mitochondria, and cytoplasmic microtubules all surrounded by a plasma membrane. The rod IS microtubular network is proposed to nucleate from the basal body (BB) (Nemet et al., 2015; May-Simera et al., 2017), which is a region of the apical IS composed of 2 centrioles, one of which - the mother centriole - is continuous with the axoneme of the CC. Cytoplasmic dynein 1 has been demonstrated to be essential for rod health as the putative motor protein complex for intracytoplasmic movement toward the minus end of microtubules in the IS (Tai et al., 1999; Insinna et al., 2010; Dahl et al., 2021b; a). The BB is also the site of the distal appendages (DAPs), which are nine radially symmetrical pinwheel-shaped blades that link the mother centriole to the plasma membrane and may serve as a gate or barrier structure at the critical IS/CC interface (reviewed in Wensel et al., 2021). The mouse rod IS also contains a ciliary rootlet, which is a filamentous cytoskeletal element that is linked to the BB and extends to the rod presynaptic terminal (Spira and Milman, 1979).

In the IS of frog rod photoreceptors, Rho has been localized with immunoelectron microscopy (immuno-EM) with the Golgi, in post-Golgi Rho-carrier vesicles, and sporadically at the IS plasma membrane (Peters et al., 1983; Papermaster et al., 1985, 1986). In early immuno-EM mammalian retina studies, Rho was localized to the IS plasma membrane in both mouse and cow rods (Jan and Revel, 1974), and in immunolabeled thin sections of pig retina, Rho was grossly localized to the IS plasma membrane, Golgi, and ONL. (Röhlich et al., 1989). More recent images of post-embedding immuno-EM or cryo-immuno-EM localization of Rho in adult mouse rods have shown minimal Rho labeling in the IS (Liu et al., 1999; Tai et al., 1999; Wolfrum and Schmitt, 2000; Hagstrom et al., 2001; Burgoyne et al., 2015; Guo et al., 2022; Chadha et al., 2019). Despite the lack of comprehensive Rho IS localization details from these studies, Rho has been consistently localized at mouse CC plasma membrane in mouse rods (Wolfrum and Schmitt, 2000; Hagstrom et al., 2001; Burgoyne et al., 2015; Chadha et al., 2019). In fluorescence localization studies using mammalian retina, including in Rho-GFP fusion knockin mouse retinas, it has been challenging to distinguish the small population of Rho molecules in the IS compared to the overwhelming amount of Rho in OS discs (Chan et al., 2004).

The network of small GTPase protein regulators of the Rho secretory pathway in the IS has been systematically defined in frog rods. In this pathway, Arf4 GTPase binds to Rho in the *trans*-Golgi network via the Rho C-terminal VxPx motif (Deretic et al., 2005), a sequence that was shown to be essential for OS targeting in frogs and zebrafish (Tam et al., 2000; Perkins et al., 2002). ASAP1, the Arf GTPase activating protein (GAP), and small GTPases FIP3 and Rab11a are then recruited to the complex to form post-Golgi Rho carrier vesicles (Mazelova et al., 2009a; Wang et al., 2012), which are then targeted to the plasma membrane by Rab8a GTPase and Rabin8, the Rab8 guanine nucleotide exchange factor (GEF) (Deretic et al., 1995; Moritz et al., 2001; Nachury et al., 2007). In mice, Arf4, Rab8 and Rab11a are not essential for Rho trafficking (Ying et al., 2016; Pearring et al., 2017) indicating the existence of redundant regulators or compensatory trafficking networks in mouse rods; however, Rab11a was shown to form protein-protein interactions with the C-terminal domain of Rho in one mouse study (Reish et al., 2014).

The mechanism for Rho vesicle docking to the IS plasma membrane in frog rods requires the SNARE (soluble N-ethylmaleimide-sensitive factor attachment protein receptor) proteins syntaxin 3 (STX3) and SNAP25 (Mazelova et al., 2009b). In the retinas from rod-specific STX3 knockout mice, Rho and the OS disc rim protein peripherin-2 are mislocalized to the IS and ONL (Kakakhel et al., 2020) demonstrating that the functional role for STX3 is maintained in mice. Furthermore, STX3 was also shown to form in vivo protein interactions with Rho (Zulliger et al., 2015). After SNARE-mediated docking, Rho-containing vesicles putatively fuse with the plasma membrane and Rho protein gets inserted into the membrane, an event that is classically mapped near the BB and CC in the apical IS of frog rods (Papermaster et al., 1985).

Overall, considering the current uncertainty regarding the impact of protein regulators of Rho biosynthesis and trafficking in mice, the complete route of rhodopsin trafficking within the IS remains largely uncharacterized despite the importance of trafficking within this critical subcellular compartment in mouse rods. In this study we used super-resolution fluorescence microscopy to perform a quantitative spatial analysis of Rho localization in mouse rods ISs. Super-resolution modalities are powerful tools for testing the subcellular localization of protein targets in mouse rod domains, such as the IS. Both structured illumination microscopy (SIM), and stochastic optical reconstruction microscopy (STORM), a single molecule localization microscopy (SMLM), were previously used to localize ciliary proteins to the nanometer-scale subcompartments of the CC in mouse rods (Robichaux et al., 2019; Potter et al., 2021). More recently, an alternative 3D STORM mode, named rapid imaging of tissues at the nanoscale STORM (RAIN-STORM), was developed to localize proteins in rod photoreceptor presynaptic terminals and post-synaptic dendrites within the mouse OPL (Albrecht et al., 2022). To enable super-resolution localization of Rho in the mouse IS, we applied techniques to reliably immunolabel Rho in the IS of mouse retina, including an OS peeling approach and nanobody targeting of Rho-GFP in Rho-GFP/+ knockin retinas, for SIM and STORM followed by a quantitative localization analysis.

## RESULTS

### Rhodopsin immunofluorescence labeling strategies in mouse retina

Previous studies established that immunolabeling the N-terminus or C-terminus of Rho labels different fractions of Rho molecules in rods from mammalian retinal tissue preparations (Hicks and Molday, 1986; Röhlich et al., 1989). The immunofluorescence staining pattern of Rho was tested using the following antibodies: 1) the 1D4 monoclonal antibody (hereafter Rho-C-1D4), which targets the last 8 amino acids in C-terminus/cytoplasmic tail of Rho, i.e. the 1D4 sequence (Molday and MacKenzie, 1983), 2) the N-terminus targeting 4D2 monoclonal antibody (Rho-N-4D2) (Hicks and Molday, 1986), and 3) a N-terminus targeting Rho-N-GTX polyclonal antibody. For immunofluorescence the Rho antibodies were first tested in immunolabeled mouse retinal cryosections imaged with confocal scanning microscopy. With this method, all the Rho antibodies targeted the OS layer almost exclusively with strong fluorescence labeling (Fig. 1 A). Rho immunofluorescence localization in mouse retinas with SIM and STORM was also tested, which required a whole-retina immunolabeling approach modified from previous STORM imaging studies (Robichaux et al., 2019; Potter et al., 2021) for both improved labeling density and thin resin sectioning of stained retinas (see Methods for details). By whole-retina labeling and SIM, the Rho-C-1D4 antibody only labeled the distal rod OS tips with fluorescence, while both N-terminal antibodies, Rho-N-4D2 and Rho-N-GTX, predominantly labeled the base of the OS where new discs are formed (Fig. 1 B). N-terminal Rho immunolabeling at the OS base is consistent with the N-termini of Rho in newly-formed evaginating discs being exposed to the extracellular space (Ding et al., 2015), while Rho-C-1D4 labeling was restricted to the OS tips. In each case, Rho protein in the IS was undetectable with immunofluorescence.

**Figure 1.**
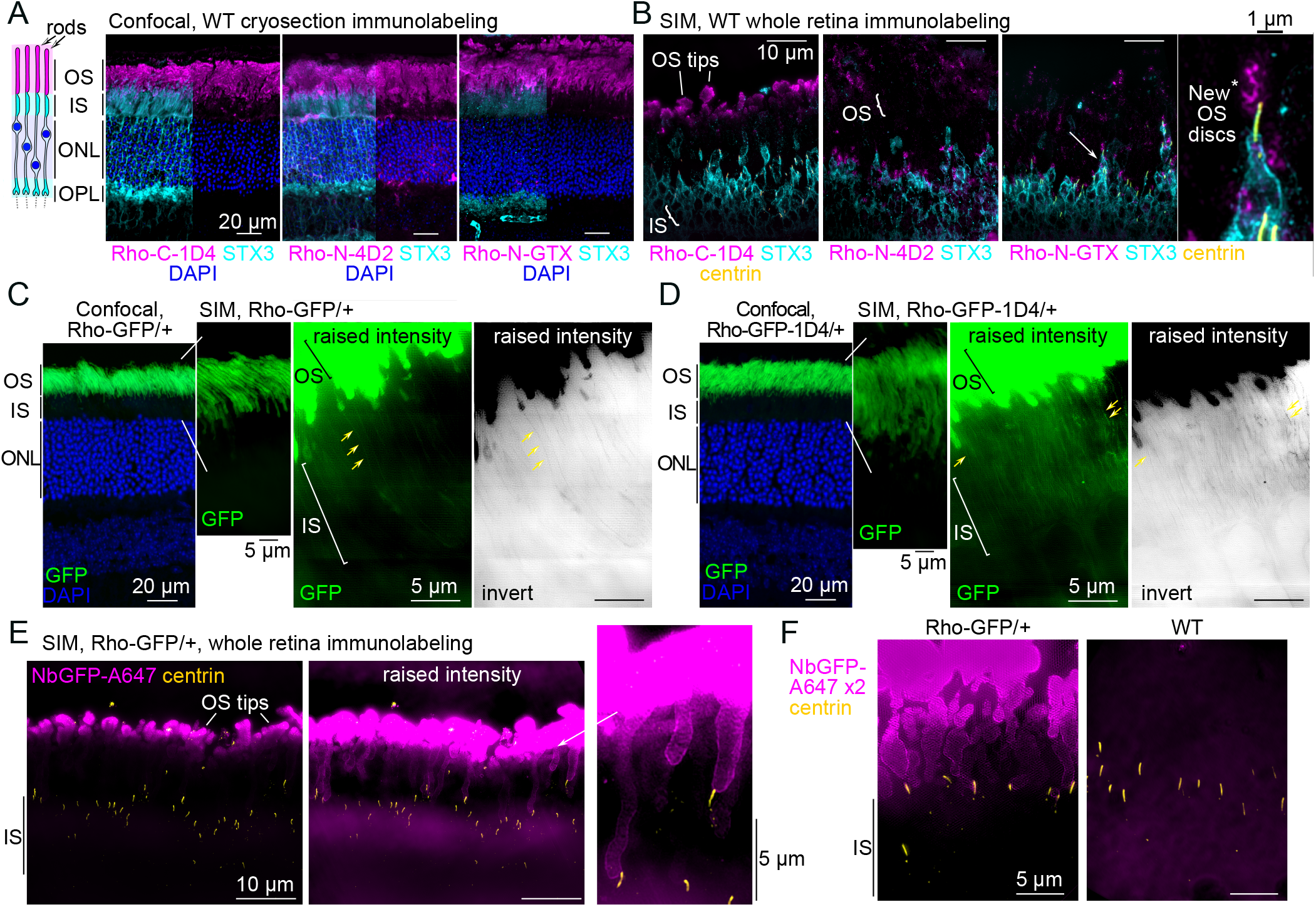
Mouse rod inner segments are non-permissive to conventional rhodopsin labeling strategies. (A) Diagram of mouse rod photoreceptor layers adjacent to z-projection confocal fluorescent images of WT mouse retinal cryosections immunolabeled for Rho (magenta), STX3 (cyan) – which labels the IS and OPL - and DAPI nuclear staining (blue) in the ONL. Rho prominently labels rod outer segments (OS). The STX3 channel is removed from part of the images for clarity. (B) Z-projection SIM images of thin (1 µm) plastic sections of WT retinas stained for immunofluorescence as whole retinas. The white arrow indicates the rod that is magnified in the Rho-N-GTX example, and the white asterisk indicates Rho-N-GTX labeling of the new OS discs in the magnified rod. Centrin (yellow) immunolabeling was used to localize the rod CC and BB. (C) Z-projection confocal image of an adult Rho-GFP/+ cryosection adjacent to z-projection SIM images of 2 µm Rho-GFP/+ cryosections. (D) Z-projection confocal image of a Rho-GFP-1D4/+ cryosection and z-projection SIM images of 2 µm Rho-GFP/+ cryosections. In both (C) and (D), a magnified region of a SIM image is shown with raised contrast and brightness (intensity) levels to depict faint IS GFP fluorescence in both heterozygous knockin mouse lines (yellow). An inverted image is shown to highlight this pattern. (E) Z-projection SIM images of Rho-GFP/+ thin plastic retina sections that were immunolabeled for NbGFP-A647 (magenta) and centrin (yellow) as whole retinas. Raised intensity images are shown to depict less intense NbGFP-A647 labeling in some proximal OS and surrounding the CC. (F) SIM images of Rho-GFP/+ and WT control retina thin sections stained as in (E) but with twice the amount of NbGFP-A647 (x2). Lack of staining in WT retinas demonstrates NbGFP-A647 specificity.

Next, two knockin Rho fusion mouse lines were used: Rho-GFP and Rho-GFP-1D4. In each knockin, GFP (green fluorescent protein) is fused to human Rho directly after the C-terminal 1D4 sequence (Chan et al., 2004). In the Rho-GFP-1D4 version of the knockin, an additional 1D4 sequence, which contains the VxPx targeting motif, is appended to the C-terminus of GFP (Robichaux et al., 2022). Although both knockin mice, as heterozygotes, have normal photoreceptor morphology and Rho OS localization, the addition of the extra 1D4 sequence was previously shown to be essential for GFP and Rho-Dendra2 OS targeting in frog and fish retinas (Tam et al., 2000; Perkins et al., 2002; Lodowski et al., 2013), and so the Rho-GFP-1D4 knockin mice was included in this mouse study to account for any possible subcellular localization requirement of the 1D4 sequence.

In confocal images of Rho-GFP/+ and Rho-GFP-1D4/+ retinas, GFP fluorescence was confined to in the OS layer, and only a faint, noise-like GFP signal was visualized in the IS layer in thin (2 µm) retinal cryosections from both knockin mice using SIM (Fig. 1 C, D). Therefore, a GFP-specific nanobody was obtained, purified, and conjugated to Alexa 647, a bright and stable fluorophore compatible with both SIM and STORM, to both enhance the GFP signal in our knockin mouse retinas and to overcome the issue of GFP bleaching during our whole retina processing technique. Nanobodies are small (∼15 kDa) single chain camelid immunoglobulins that are ideal for super-resolution fluorescence microscopy (Ries et al., 2012). The specificity of the GFP nanobody Alexa 647 conjugate (abbreviated NbGFP-A647) was validated with western blotting and confocal immunofluorescence (Fig. S1 A-C), in which WT retinas were used as controls for Rho-GFP labeling with NbGFP-A647. In each test, NbGFP-A647 properly labeled Rho-GFP or Rho-GFP-1D4 with no evidence of non-specific labeling in WT controls. Notably, the Rho-C-1D4 antibody properly labeled the Rho-GFP-1D4 protein band in western blots of Rho-GFP-1D4/+ retinal lysates (Fig. S1 B). To validate these mouse lines further, we used a deglycosylation assay to confirm that Rho-GFP and Rho-GFP-1D4 fusion proteins, from Rho-GFP/+ and Rho-GFP-1D4/+ retina lysates respectively, were properly glycosylated like the endogenous mouse Rho protein from the same lysates (Fig. S1 D, E).

Next, whole retina immunolabeling of Rho-GFP/+ and WT retinas with NbGFP-A647 was performed. In SIM images of thin sections from these Rho-GFP/+ retinas, NbGFP-A647 fluorescence was again predominantly limited to the distal OS tips, and only a few rods had any Rho-GFP labeling in the proximal OS near the centrin-positive CC (Fig. 1 E). Immunolabeling of centrins, which are calcium-binding proteins localized specifically to the CC and BB centrioles in mouse rods (Trojan et al., 2008; Robichaux et al., 2019), was used to label the positions of the CC and BB as the apical edge of the IS throughout the study. Even when we doubled the amount of NbGFP-A647 used during labeling, the Rho-GFP-localized fluorescence was still predominantly found in distal OS tips of Rho-GFP/+ retinas with no detectable fluorescent signal in the IS (Fig. 1 F).

### Outer segment peeling of mouse retinas enables immunofluorescence detection of rhodopsin in the photoreceptor inner segment layer

To improve the penetration of our immunoreagents to the rod IS, a retinal peeling technique was used in which mouse retinas were iteratively adhered to and removed from filter paper to physically detach OSs from the retina in order to generate IS-enriched retina samples (Rose et al., 2017, 2021) (Fig. 2 A). OS removal using between 4 and 8 paper peeling cycles was tested using standard transmission electron microscopy (TEM). 8 peeling cycles resulted in increased OS removal and so this number was used throughout this investigation to generate IS-enriched retinas. (Fig. S2 A). After generating IS-enriched retinas from Rho-GFP/+ mice, the enrichment of IS proteins was validated with western blotting, where the amount of OS proteins, Rho-GFP, endogenous mouse Rho (labeled with Rho-C-1D4 antibody), and ROM1, were reduced compared to full retina control samples, while the levels of IS proteins STX3 and Rab11a were unchanged between conditions (Fig. 2 B).

**Figure 2.**
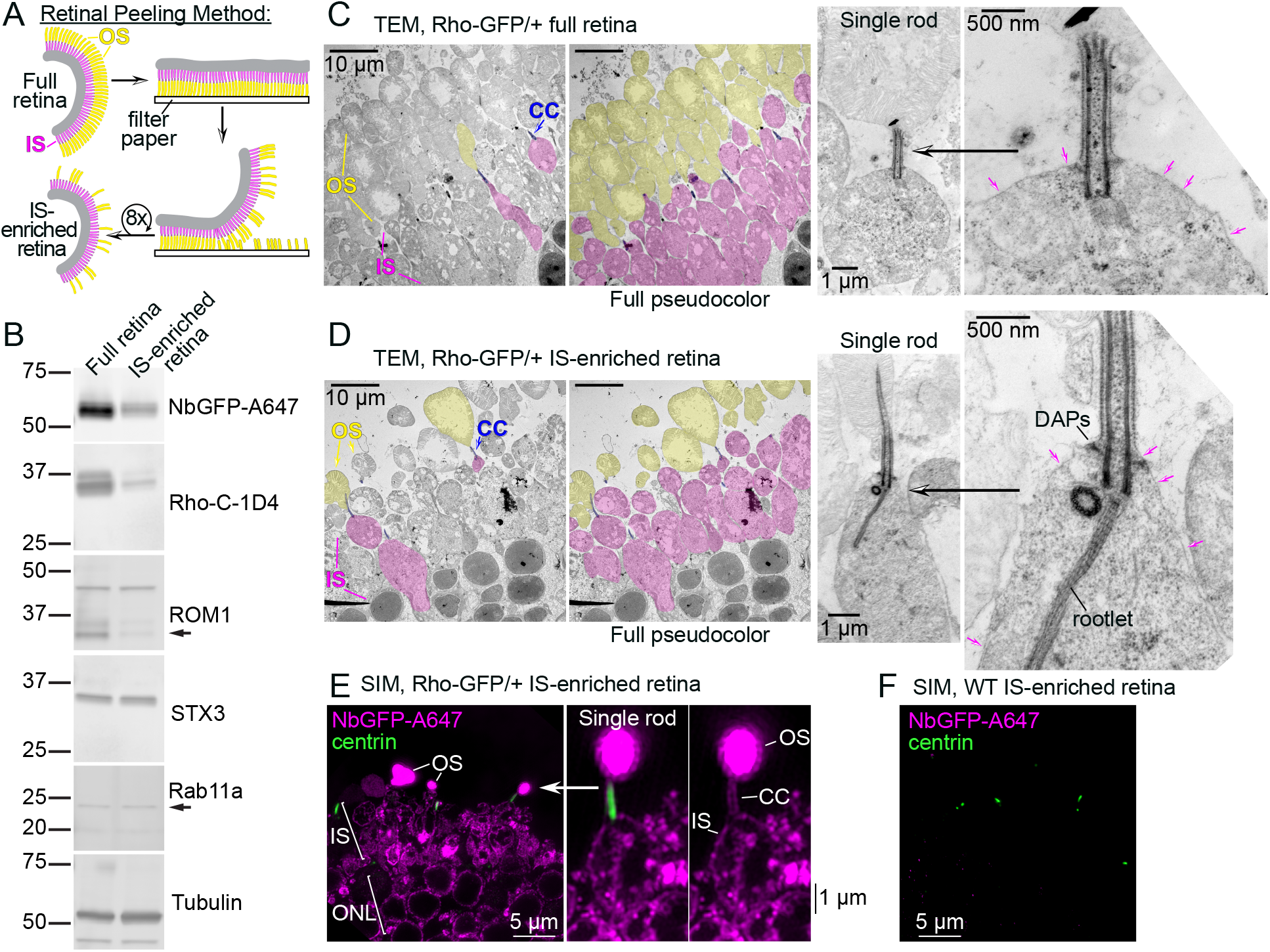
Outer segment peeling generates inner segment-enriched mouse retina samples. (A) Diagram of the mouse retina OS peeling method. Mouse retina slices are iteratively peeled from filter paper 8 times to remove rod outer segments (OS, yellow) from the inner segment (IS) layer (magenta). (B) Western blot analysis comparing control full retina samples vs. retinas after OS peeling/IS-enriched retinas; both from Rho-GFP/+ mice. 2% of total volume from lysates from 1 retina (either control or peeled) were used from each condition for SDS-PAGE. Molecular weight marker sizes are indicated in kDa. Immunolabeled bands from antibodies targeting OS proteins, including NbGFP-A647, Rho-C-1D4 and ROM1 (black arrow points to the ROM1 monomer band), were reduced in peeled lysates, whereas IS proteins STX3 and Rab11a were roughly equal demonstrating IS enrichment. Tubulin immunoblotting was performed as a loading control. (C) Control full retina slices and (D) IS-enriched retinal slices from Rho-GFP/+ mice were fixed and stained for resin embedding, ultrathin sectioning and TEM ultrastructure analysis. TEM images were pseudocolored to point out key rod structures as follows: OS = yellow, IS = magenta, connecting cilia (CC)/basal bodies = blue. Single rod examples are magnified from both conditions to emphasize intact CC ultrastructure and the preservation of the IS plasma membrane (magenta arrows). The locations of the distal appendages (DAPs) and rootlet are annotated in (D). (E) Z-projection SIM images of Rho-GFP/+ IS-enriched retinas immunolabeled with NbGFP-A647 (magenta) and centrin antibody (green). Fluorescence in the single rod example corresponding to the OS, IS and CC are annotated. (F) Control SIM image of a WT IS-enriched retina immunolabeled and imaged as in (E) to demonstrate NbGFP-A647 labeling specificity. Scalebar values match adjacent panels when not labeled.

Next, Rho-GFP/+ retinas were fixed for TEM immediately after peeling/IS-enrichment to evaluate if the IS and CC ultrastructure were maintained after OS removal. Compared to the overall morphology of unpeeled Rho-GFP/+ rod photoreceptors (Fig. 2 C), IS-enriched retinas contained fewer OS with a generally intact IS layer (Fig. 2 D). In higher magnification TEM micrographs, IS-enriched rod examples had preserved CC structure that were typically attached to a remaining OS, and these rods had an intact IS plasma membrane and cytoplasm (Fig. 2 D, Fig. S2 B)

IS-enriched retinas from Rho-GFP/+ mice were then immunolabeled with NbGFP-A647 using whole retina immunolabeling and SIM as before. In this case, presumably due to the removal of many of the OSs in these retinas that would otherwise soak up all available immunoreagents, NbGFP-A647 fluorescence was now observed in the rod CC, IS and ONL regions among the sparse remaining OSs in these sections (Fig. 2 E). WT IS-enriched retinas were also stained with NbGFP-A647 to demonstrate labeling specificity (Fig. 2 F).

### Rhodopsin is localized at the mouse rod inner segment plasma membrane

Compared to confocal images of Rho-GFP/+ IS-enriched retinas co-immunolabeled with NbGFP-A647 and centrin antibody (Fig. 3 A), SIM clearly improved the signal-to-noise ratio of the immunofluorescence in our retina sections, which was further enhanced with 3D deconvolution (Fig. 3 B). In SIM images of single Rho-GFP/+ rods, Rho-GFP was prominently localized at the apparent boundary of the IS in a continuous string of fluorescence that was contiguous with the CC plasma membrane (Fig. 3 B). Bright puncta of Rho-GFP were also observed in the proximal IS region, known as the myoid (Golgi-rich) IS (Fig. 3 B, asterisks); however, in single rod views, internal Rho-GFP in the distal/ellipsoid IS cell body was not reliably observed in any particular subcompartment. STORM single molecule localization of NbGFP-A647 in Rho-GFP/+ IS-enriched retinas demonstrated the same localization pattern of Rho-GFP at the apparent rod IS plasma membrane (Fig. 3 C).

**Figure 3.**
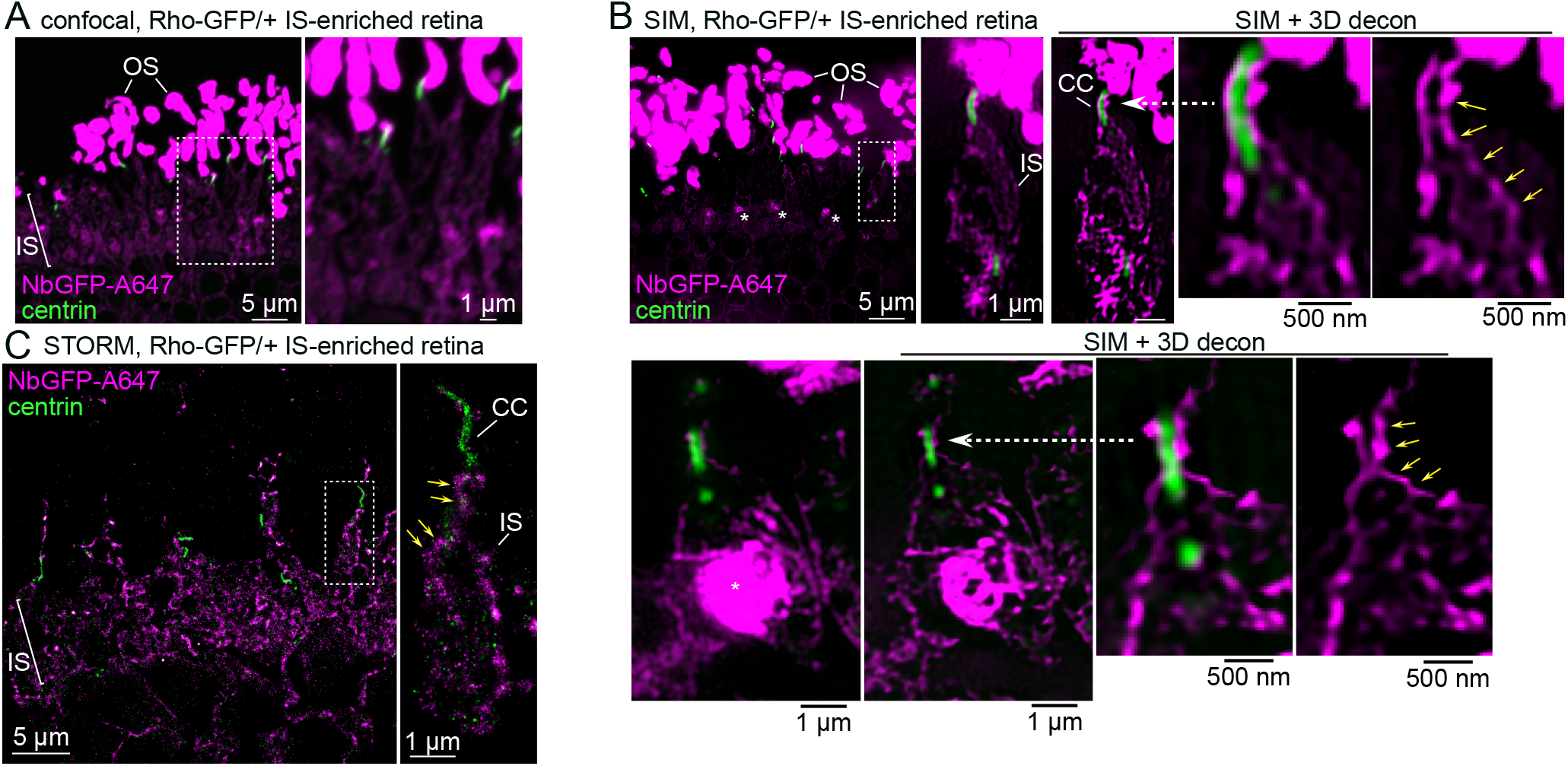
A fraction of rhodopsin-GFP molecules are localized at the rod inner segment plasma membrane. (A) Confocal z-projection and (B) SIM z-projection images of an IS-enriched, Rho-GFP/+ retina section immunolabeled with NbGFP-A647 (magenta) and centrin antibody (green). Single rod examples are magnified in (B) to highlight Rho-GFP localization along the boundaries of the inner segment (IS) and connecting cilium (CC). This same SIM single rod images are also shown after 3D-deconvolution processing (SIM + 3D decon). Yellow arrows indicate a continuous line of Rho-GFP fluorescence between the IS and CC. Asterisks indicate strong Rho-GFP puncta staining in the proximal IS/myoid region. (C) STORM reconstruction of an IS-enriched Rho-GFP/+ retina section. In the magnified single rod examples, yellow arrows indicate Rho-GFP molecules located at the IS boundary. Magnified regions are indicated throughout with either a dashed box or a dashed white arrow. Some magnified images are rotated so that the OS end of the rod is at the top of the image. Scalebar values match adjacent panels when not labeled.

We sought a reliable antibody marker of the IS plasma membrane so that single ISs could be identified in our localization experiments. Antibodies that target syntaxin 3 (STX3), a t-SNARE protein previously localized at the IS plasma membrane in mouse retina (Chuang et al., 2007; Datta et al., 2015; Kakakhel et al., 2020), and the Na/K-ATPase channel, which has also been used as a general IS marker (Kwok et al., 2008; Dilan et al., 2019), were tested. Using SIM and STORM super-resolution microscopy experiments, the STX3 antibody resulted in more densely labeled IS boundaries of individual rods (Fig. 4 A, B). Therefore, STX3 was used as the IS plasma membrane marker in subsequent experiments.

**Figure 4.**
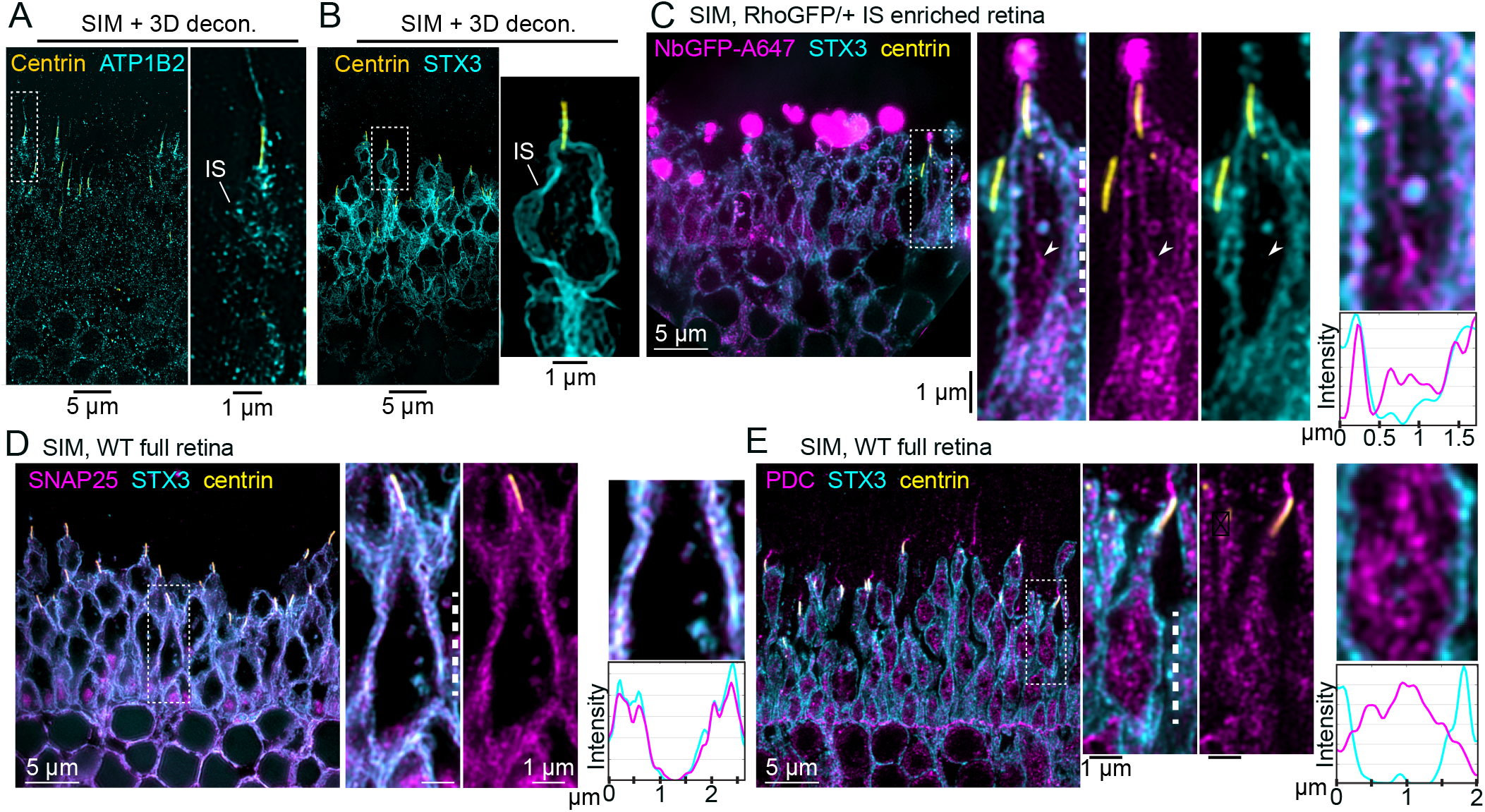
Rhodopsin colocalized with syntaxin 3 at the rod inner segment plasma membrane. (A) Example SIM z-projection image (processed with 3D deconvolution), of a thin plastic section from a WT mouse retina immunolabeled with antibodies for anti-centrin (yellow), to label the CC and BB, and anti-Na,K ATPase B2 subunit (ATP1B2, cyan) to label the IS plasma membrane. (B) Example SIM z-projection image with 3D deconvolution of a thin plastic section from a mouse retina immunolabeled with anti-centrin and anti-STX3 (cyan) to label the IS plasma membrane. (C) SIM image of an IS-enriched Rho-GFP/+ retina immunolabeled for NbGFP-A647 (magenta), syntaxin 3 (STX3), and centrin (yellow). In the magnified single rod example, an arrowhead indicates a cytoplasmic IS region with Rho-GFP fluorescence and no STX3 fluorescence. To demonstrate Rho-GFP colocalization with STX3 at the IS plasma membrane, a row average intensity plot is shown for a portion of the IS from the magnified single rod example marked with a dashed line. SIM images of (D) SNAP25 (magenta) and (E) phosducin (PDC, magenta) immunolabeling in WT full retina sections that are each co-labeled with STX3 (cyan) and centrin (yellow) antibodies. For both magnified single rod examples, row intensity plots are provided for the IS regions marked by white dashed lines. Fluorescence intensities are normalized on all plots for clarity. Throughout the figure, magnified regions are indicated with a dashed box. Some magnified images are rotated so that the OS end of the rod is at the top of the image. Scalebar values match adjacent panels when not labeled.

To test if Rho-GFP localizes at the rod IS plasma membrane, IS-enriched Rho-GFP/+ retinas were co-immunolabeled for NbGFP-A647 along with the STX3 antibody to mark the location of the IS plasma membrane. In SIM images, RhoGFP + STX3 fluorescent signals were partially overlapped in the rod IS, including a near co-localization at the plasma membrane (Fig. 4 C), a result that was recapitulated for Rho-GFP-1D4 (Fig. S2 C), confirming Rho-GFP/Rho-GFP-1D4 localization at the IS plasma membrane. In both cases, internal Rho-GFP and Rho-GFP-1D4 did not completely co-localize with STX3. As a positive control for IS plasma membrane colocalization, immunolabeling of SNAP25, another IS SNARE complex protein like STX3 (Mazelova et al., 2009b; Kakakhel et al., 2020), was used in full WT retinas for SIM. In single rods from these retinas, STX3 and SNAP25 completely colocalized at the plasma membrane (Fig. 4 D). As a cytoplasmic protein control, immunolabeling of phosducin (PDC), a soluble IS chaperone protein (Sokolov et al., 2004), was used in WT retinas for SIM. In this case, PDC within single rod ISs was completely internal, corresponding to its cytoplasmic localization, and did not colocalize with STX3 at the plasma membrane (Fig. 4 E). Importantly, the correct and unperturbed localization of SNAP25, PDC and STX3 in these data demonstrates the cellular preservation of the rod IS in retinas that are processed for immunofluorescence for either SIM or STORM.

Although NbGFP-A647 in Rho-GFP knockin mice provides specific and rigorous Rho labeling, the endogenous mouse Rho localization pattern was also tested in the ISs of WT mouse rods. In IS-enriched WT retinas that were immunolabeled with Rho-C-1D4 antibody, the endogenous Rho localization pattern matched the Rho-GFP IS pattern, including localization at the IS plasma membrane and CC membrane, in addition to prominent bright puncta labeling in the myoid IS (Fig 5 A). Mouse retinal immunolabeling for the cis-Golgi marker GM130 (Nakamura et al., 1995) was previously shown to label a bright, amorphous punctum in the IS myoid (Sedmak and Wolfrum, 2010; Grossman et al., 2011; Pearring et al., 2017), a localization pattern that was recapitulated here with SIM (Fig. 5 B). IS-enriched Rho-GFP/+ retinas were co-immunolabeled with NbGFP-A647, GM130, STX3 and centrin antibodies to attempt to co-localize the bright myoid Rho puncta with GM130. However, in SIM images from these retinas, the Rho-GFP puncta did not directly colocalize with GM130. Instead, the Rho-GFP fluorescence closely surrounded the GM130+ cis-Golgi puncta in a reticulated fashion (Fig. 5 C).

**Figure 5.**
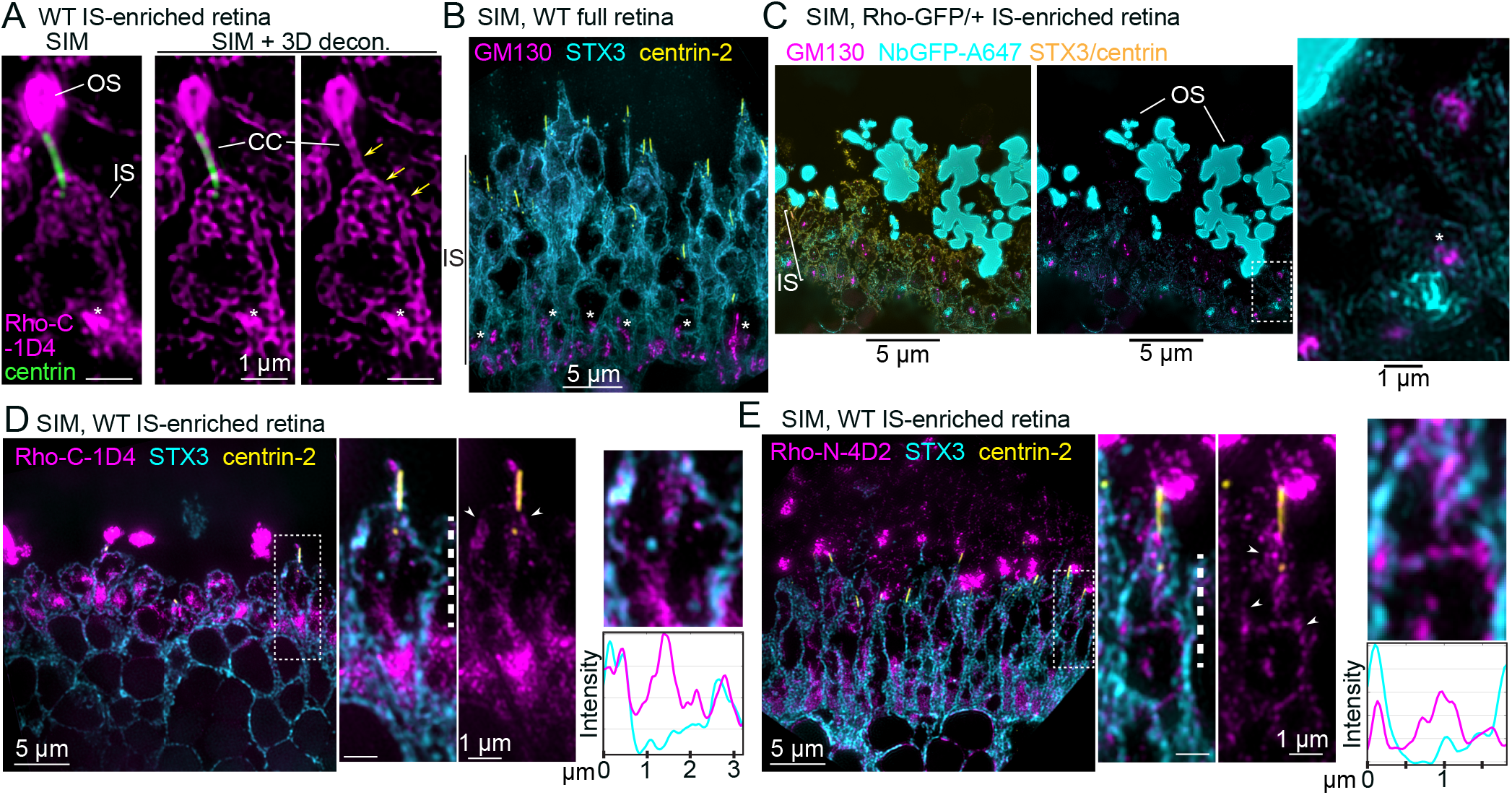
Endogenous inner segment mouse rhodopsin is located near the cis-Golgi and at the plasma membrane. (A) SIM z-projection images of a single rod example from an IS-enriched, WT retina section immunolabeled with Rho-C-1D4 (magenta) and centrin antibody (green). Rho-C-1D4 is localized along the boundaries of the IS and CC (yellow arrows). The OS is indicated. (B) SIM z-projection image of a WT full retina section immunolabeled with GM130 (magenta), STX3 (cyan) and centrin-2 (yellow) antibodies. Asterisks indicate strong GM130 puncta staining in the proximal IS/myoid region corresponding to the cis-Golgi in each rod IS. (C) SIM z-projection image of a Rho-GFP IS-enriched retina section immunolabeled with NbGFP-A647 (cyan), along with GM130 (magenta), STX3 (yellow) and centrin (yellow) antibodies. The white arrow indicates a magnified portion of the IS layer, in which an asterisk indicates a GM130 punctum located directly near a Rho-GFP punctum in the myoid of a rod. (D, E) SIM images of an WT IS-enriched retinas immunolabeled with either (D) Rho-C-1D4 antibody (magenta) or (E) Rho-N-4D2 antibody (magenta); both are co-labeled with STX3 (cyan), and centrin-2 (yellow) antibodies. Partial endogenous Rho colocalization with STX3 at the IS plasma membrane is indicated with white arrowheads, and row average intensity plots are from the magnified portion of the IS corresponding to the region in single rod example images marked with a dashed line. Fluorescence intensities are normalized on both plots for clarity. Throughout the figure, magnified regions are indicated with either a dashed box or a dashed white arrow. Some magnified images are rotated so that the OS end of the rod is at the top of the image. Scalebar values match adjacent panels when not labeled.

Next, WT IS-enriched retinas were immunolabeled with either Rho-C-1D4 or Rho-N-4D2 along with STX3 to confirm endogenous mouse Rho localization at the IS plasma membrane with SIM. In each case, Rho colocalization with STX3 at the IS PM was observed along with a discontinuous internal labeling in the IS that was dense in the myoid IS (Fig. 4 D, E and Fig. S2 D, E). Overall the localization of endogenous mouse Rho in the IS closely matched the Rho-GFP and Rho-GFP-1D4 IS localization pattern in Rho-GFP/+ and Rho-GFP-1D4 rods, respectively.

### STORM spatial analysis of Rho subcellular localization in mouse rod inner segments

STORM was used to generate single molecule reconstructions of Rho localization in the IS from IS-enriched retinal samples. Molecule coordinates from STORM reconstructions of rod ISs were extracted to quantitatively test if a significant fraction of Rho molecules in the IS localized at the IS plasma membrane using a spatial localization analysis to measure the distance of Rho STORM molecules to the IS boundary or “hull”. In all STORM data, reconstructed molecules from STX3 immunolabeling were used to manually define the rod IS hull, and centrin-2 was used as a marker of the CC and BB. Only rod ISs that contained a centrin-2 positive CC/BB were included in the analysis for consistency. The results, as detailed with statistical analysis below, strongly support the conclusion that Rho-containing internal membranes are preferentially localized near the IS plasma membrane.

First, STORM reconstructions from Rho-GFP/+ IS-enriched retinal sections immunolabeled with NbGFP-A647 were analyzed and single rod ISs were identified. STORM molecules within the STX3+ IS hull were isolated from the STORM reconstruction of the retina section and plotted separately (Fig. 6 A, Fig. S3 A). In isolated STORM plots, Rho-GFP molecules partially overlapped with STX3 molecules in most examples, including at the IS hull. For comparison, a random distribution of molecules was plotted within the same IS hulls. Based on these plots, distance-to-hull values, which are the nearest distance of each molecule to the IS hull, were calculated for the Rho-GFP, STX3, and the random STORM molecules from each rod IS. Based on these data, a large fraction of Rho-GFP molecules were located within 0.2 µm of the IS hull, with distributions typically matching STX3 molecules as opposed to random molecule distributions (Fig. 6 A, Fig. S3 A). In alternate rod examples where Rho molecules are more densely localized internally, the distribution of Rho-GFP distance-to-hull values shifted to higher mean distances (Fig S3 A); however, Rho-GFP reconstructed molecules from these STORM plots are still consistently localized along the IS hull. The STORM analysis was repeated in Rho-GFP-1D4/+ retinas using NbGFP-A647 labeling, and the distributions of Rho-GFP-1D4 distance-to-hull values were also consistently weighted toward lower mean distances (Fig. S3 B).

**Figure 6.**
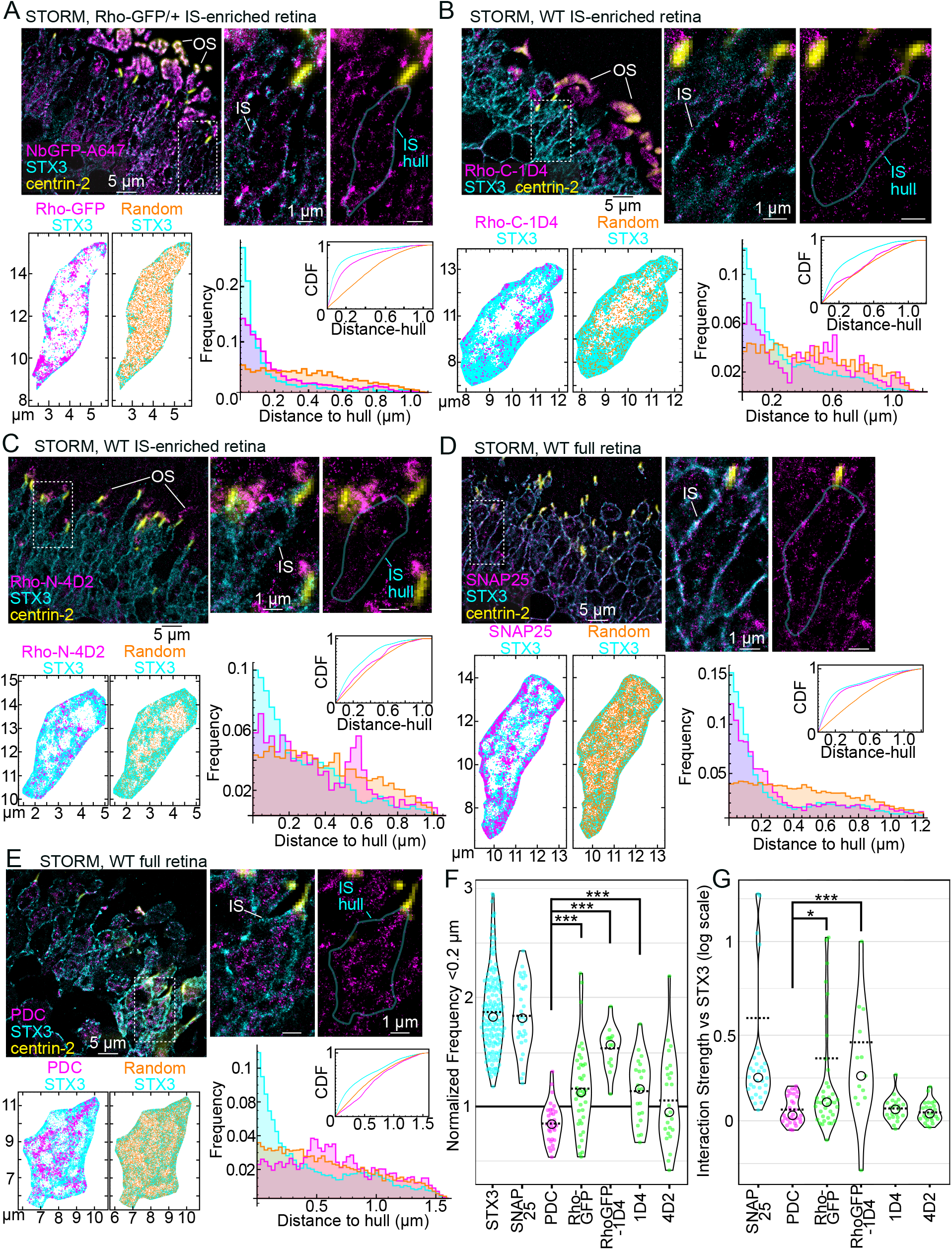
STORM spatial analysis of rhodopsin localization in rod inner segments. (A) STORM reconstruction of an IS-enriched Rho-GFP/+ retina immunolabeled with NbGFP-A647 (magenta) and STX3 (cyan) and centrin-2 (yellow) antibodies. Outer segments (OS) are indicated. The centrin-2+ widefield images are superimposed on STORM reconstruction images to mark the positions of rod connecting cilia and basal bodies. In single rod examples, the IS region is indicated, and the IS hull – the manually defined STX3+ IS boundary – is outlined in cyan in a duplicate image with the STX3 STORM reconstruction removed. From the rod example in (A), the Rho-GFP and STX3 STORM molecule coordinates within the IS hull are plotted (Rho-GFP molecules = 4,239, STX3 molecules = 5,215). In the adjacent plot, a random distribution of coordinates within the IS hull matching the number of Rho-GFP molecules (4,239) are plotted in orange with the STX3 molecules. Nearest distance to hull measurements for Rho-GFP, STX3 and random molecules are plotted in a frequency graph and a cumulative distribution function (CDF) graph. Colors in the graphs match the molecule plots. (B-E) STORM reconstructions of IS-enriched WT retinas immunolabeled with either Rho-C-1D4 antibody (B) or Rho-N-4D2 (C), and full WT retinas immunolabeled with (D) SNAP25 or (E) PDC; all are co-immunolabeled with STX3 and centrin-2 antibodies. Single rod examples and IS hulls are indicated. Molecules coordinates are plotted along with corresponding randomly plotted molecules (orange) as in (A), and distance to hull measurements are plotted as frequency and CDF graphs. Molecule counts: (B) Rho-C-1D4 = 1,430, STX3 = 24,596, Random = 1,430; (C) Rho-N-4D2 = 3,199, STX3 = 24,044, Random = 3,199; (D) SNAP25 = 7,871, STX3 = 28,817, Random = 7,871; (E) PDC = 14,556, STX3 = 34,544, Random = 14,556. Dashed boxes indicate the single rod examples in magnified images. Scalebar values match adjacent panels when not labeled. (F) For each rod analyzed with STORM, the distance to hull frequency within 0.2 µm was divided by the distance to hull frequency within 0.2 µm of the corresponding random coordinates to acquire “normalized frequency <0.2 µm” values, which are compared as violin plots. STX3 n value (for number of rods analyze) =162, SNAP25 n=31, PDC n=31, Rho-GFP n=40, Rho-GFP-1D4 n=13, Rho-C-1D4 n=21 and Rho-N-4D2 n=25 conditions were tested for statistical significance using the Mann-Whitney U test. PDC vs Rho-GFP ***P-value<0.001; PDC vs Rho-GFP-1D4 ***P-value<0.001; PDC vs 1D4 ***P<0.001. (G) The same STORM data were used to perform Mosaic interaction analyses to test the colocalization between STX3 molecules and the other immunolabeled targets from the same rod ISs. Interaction strength values are compared as violin plots on a log scale. PDC vs Rho-GFP, Rho-C-1D4 and Rho-N-4D2 (same n values as (F)) were tested for statistical significance using the Mann Whitney U test. PDC vs Rho-GFP *P=0.026; PDC vs Rho-GFP-1D4 ***P<0.001. In violin plot graphs, circles = median values and dashed lines = mean values.

Next, the same STORM spatial analysis was performed in WT IS-enriched retinas immunolabeled with Rho-C-1D4 and Rho-N-4D2 antibodies (Fig. 6 B-C, Fig. S3 C, D). In these rod reconstructions, the distribution of Rho molecule distance-to-hull measurements varied based on the internal distribution of Rho in any given rod, but as with the Rho-GFP data, endogenous Rho molecules were always localized to some degree near the IS hull, typically as a large fraction of molecules localized within 0.2 µm of the IS hull.

STORM IS molecule coordinates from SNAP25 and PDC immunolabeled WT full retina samples were also analyzed as spatial analysis controls. As with our SIM data, SNAP25 molecules co-localized with STX3 molecules in rod STORM reconstructions, and SNAP25 distance-to-hull distributions also closely matched STX3 (Fig. 6 D, Fig. S3 E). PDC IS reconstructed STORM molecules, on the other hand, were almost completely internal, and distance-to-hull measurements accumulated at greater distances from the IS hull, completely distinct from STX3 distance distributions from the same rod examples (Fig. 6 E, Fig S3 F).

All distance-to-hull measurements were aggregated from this STORM rod IS spatial analysis, including for STX3. The frequency of molecules <0.2 µm from the IS hull for any STORM target (e.g. Rho, STX3) was normalized to the corresponding random <0.2 µm frequency from the same rod reconstruction to account for different IS areas and labeling densities in our STORM data. Normalized frequency values were plotted for comparison (Fig. 6 F), and STX3 and SNAP25 normalized values were clearly greater than 1 in aggregate, confirming a greater than random localization of STX3 and SNAP25 within 0.2 µm of the IS hull. Rho-GFP, Rho-GFP-1D4, Rho-C-1D4 and Rho-N-4D2 normalized values all were, in aggregate, nearer to 1 due to the variable fraction of internal Rho IS molecules that were also measured in this analysis. However, one sample t-tests were performed for all Rho conditions to statistically compare the normalized data to 1. Based on this test, Rho-GFP, Rho-GFP-1D4 and Rho-C-1D4 data were statistically different compared to 1 (two-tailed P-values: Rho-GFP vs 1 P= 0.0117, Rho-GFP-1D4 vs 1 P< 0.0001, Rho-C-1D4 vs 1 P= 0.0349) confirming that the fraction of Rho molecules in those rods localized within 0.2 µm of the IS hull is larger than expected for randomly distributed molecules. Furthermore, by direct comparison, Rho-GFP, Rho-GFP-1D4 and Rho-C-1D4 normalized <0.2 µm frequency values were significantly greater than cytoplasmic PDC <0.2 µm frequency values (Fig. 6F). Normalized frequency values of distance-to-hull measurements within 0.1 µm and 0.3 µm were also aggregated for all conditions, and the same general distribution of the aggregated data was observed (Fig. S4 A, B). In summary, despite a variable fraction of internally localized Rho in STORM-reconstructed rods, most rod ISs contain a non-random fraction of Rho molecules that localized near the IS plasma membrane within 0.1-0.3 µm from the IS hull with a greater frequency compared with internally-localized STORM PDC molecules.

The same rod IS STORM data was used for a Mosaic spatial interaction analysis of colocalization (Shivanandan et al., 2013) comparing the spatial distribution patterns of STX3 vs. SNAP25, PDC, Rho-GFP, Rho-GFP-1D4, Rho-C-1D4, and Rho-N-4D2 within our manually defined IS hull regions. Interaction strength values for these comparisons were calculated and normalized to the interaction strength values of random molecule distributions (each compared to STX3) to account for rod to rod variations in IS area and labeling density. When aggregated, SNAP25 vs. STX3 had the greatest normalized interaction strength scores, as expected, while both Rho-GFP vs. STX3 and Rho-GFP-1D4 vs STX3 normalized interaction strength values were significantly greater than PDC vs. STX3 (Fig. 6 G), indicating a strong colocalization between Rho-GFP/Rho-GFP-1D4 and STX3 in the STORM data. Rho-C-1D4 and Rho-N-4D2 interaction strength values compared to STX3 were not statistically different from PDC likely due to larger internal Rho densities in these rods that do not colocalize with STX3.

Immuno-EM was performed to validate our STORM single molecule localization findings, using a variation of the same whole retina immunolabeling mouse retina protocol used for SIM and STORM. Using this method, Nanogold labeled STX3 was accumulated at the rod IS plasma membrane (Fig. S4 C), while Rho-C-1D4 Nanogold labeling in rods from WT IS-enriched retina also aligned to the IS plasma membrane border (Fig. S4 D). Ultrastructural damage in immuno-EM retina samples could not be avoided due to the harsh immuno-EM processing steps that have been previously described (Kellenberger et al., 1992; Humbel et al., 1998; Baschong and Stierhof, 1998); however, these EM-based localization results support our super-resolution fluorescence localization of Rho IS at the IS plasma membrane in mouse rods.

Coimmunoprecipitation (co-IP) experiments were performed using IS-enriched retina and Triton X-100 detergent extraction to identify IS Rho protein-protein interactions in vivo to support our STORM colocalization data. First, using Rho-GFP/+ IS-enriched retinal lysates, co-IPs were performed with GFP-Trap magnetic beads to capture Rho-GFP protein, which immunoprecipitated with both STX3 and SNAP25 (Fig. 7 A). GC1 (guanylate cyclase 1), a known protein interactor with Rho (Pearring et al., 2015) was the positive control, and PDC was the negative control. Similar co-IP experiments using IS-enriched Rho-GFP-1D4 retinal lysates and GFP-Trap beads yielded the same interaction results as the Rho-GFP pulldown (Fig 7 B). Next anti-1D4 IgG bound agarose beads were used to capture endogenous mouse Rho from IS-enriched Rho-GFP/+ lysates using the same co-IP controls as Fig. 7 A-B. Here, Rho-GFP protein was immunoprecipitated, demonstrating that the endogenous mouse Rho interacts with human Rho-GFP in Rho-GFP/+ rods (Fig 7 C). STX3 was also immunoprecipitated with endogenous mouse Rho. Finally, using WT IS-enriched retinas and anti-1D4 IgG beads, STX3 was immunoprecipitated with Rho (Fig. 7 D) demonstrating this protein-protein interaction *in vivo*.

**Figure 7.**
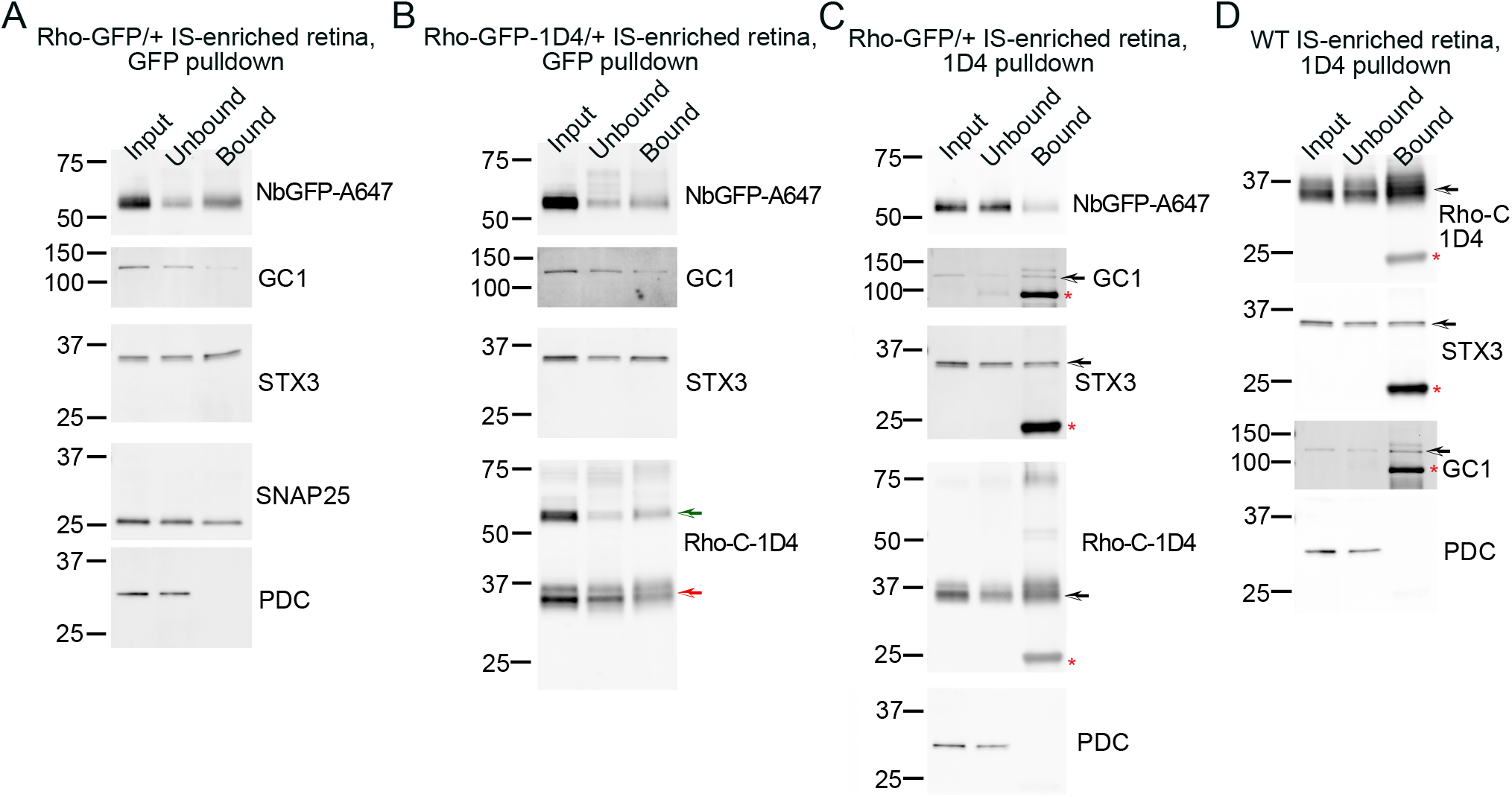
Syntaxin 3 coimmunoprecipitates with rhodopsin in vivo. Coimmunoprecipitation results from (A) Rho-GFP/+ IS-enriched retinal lysates incubated with GFP-Trap agarose beads, (B) Rho-GFP-1D4/+ IS-enriched retinal lysates incubated with GFP-Trap agarose beads, (C) Rho-GFP/+ IS-enriched retinal lysates incubated with anti-1D4 IgG agarose beads, and (D) WT IS-enriched retinal lysates incubated with anti-1D4 IgG agarose beads. In all western blots, input lanes correspond to 2% (% vol/vol) of the starting lysate volume, unbound lanes correspond 2% (% vol/vol) of lysate volume post bead incubation, and bound lanes correspond to half the total eluate from each coimmunoprecipitation. Antibodies/nanobodies used for immunodetection are listed to the right of each corresponding western blot scan. Molecular weight marker sizes are indicated in kDa. Black arrows mark the correct size bands when other bands are present on the scan. Red asterisks indicate mouse IgG bands. In (B) the red arrow indicates the endogenous Rho band and the green arrow indicates the Rho-GFP-1D4 band.

### Rhodopsin surrounds the distal appendages at the inner segment-connecting cilium junction

Since we observed a continuous string of Rho fluorescence between the IS plasma membrane and into the CC membrane in mouse rods (Fig. 3 B, Fig. 5 A), we next tested Rho molecule localization relative to the DAPs, which are the BB structures at the IS/CC interface in rods. The DAPs are 9 proteinaceous blades that project radially from the mother centriole at the base of the CC and extend to the plasma membrane (Wensel et al., 2021). In mammalian primary cilia, the DAPs were shown to function as binding sites for IFT (intraflagellar transport) proteins, which facilitate active ciliary transport, and to function in the gating of the ciliary GPCRs Smoothened and SSTR3 (Yang et al., 2018).

Here, an antibody targeting the DAP protein CEP164 was used to mark the position of the DAPs in rods, and in SIM images, the CEP164+ DAPs were localized as a well-defined line of fluorescence at the proximal end of the CC, which is the IS-CC interface (Fig. 8 A). Next, Rho-GFP/+ IS-enriched retinas were immunolabeled with NbGFP-A647 along with CEP164 and centrin antibodies to localize Rho at the DAPs using SIM and STORM. In these data, Rho-GFP molecules were again localized as continuous strings of fluorescence between the IS and CC plasma membranes but in a pattern surrounding the CEP164+ DAP blades (Fig. 8 B-C) with only a few STORM molecules co-localized with the CEP164+ DAPs. These results suggest that Rho is not integrated into the DAPs, but is likely associated with the surrounding plasma membrane between the IS and CC.

**Figure 8.**
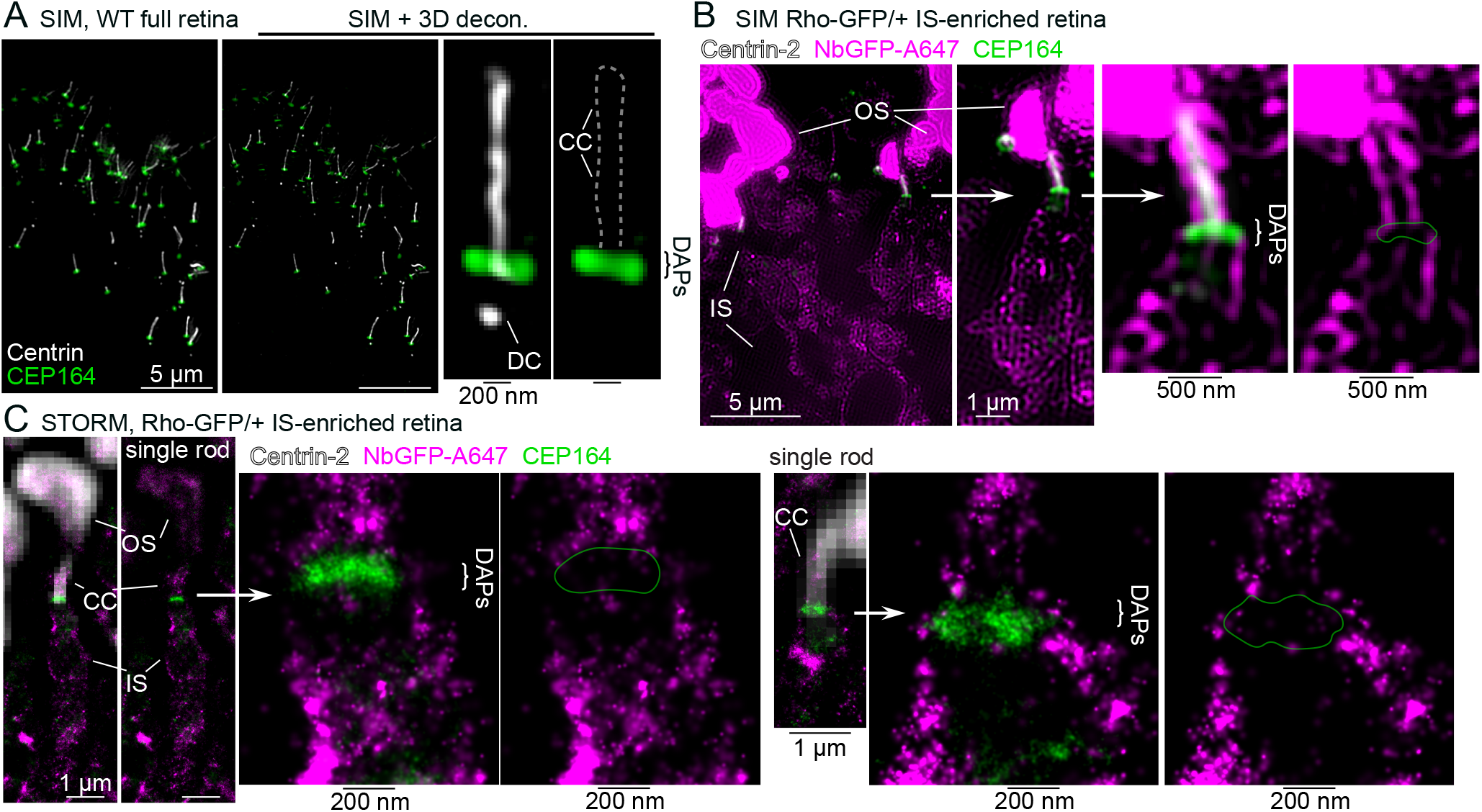
Rhodopsin localizes around the distal appendages in mouse rods. (A) SIM z-projection images of a WT full retina section immunolabeling with a centrin antibody that marks the location of rod CC and the daughter centriole (DC) of the BB in white, along with a CEP164 antibody that labels the distal appendages (DAPs) in green. The SIM retina image is also shown after 3D-deconvolution processing (SIM + 3D decon.) and a single rod cilium example is magnified. (B) SIM image from peeled Rho-GFP/+ mouse labeled with NbGFP-A647 (magenta), anti-CEP164 antibody (green) and anti-centrin antibody (white). A single rod is magnified and the DAPs region is further magnified. The remaining OSs and the IS region are indicated. (C) STORM reconstruction single rod examples and magnified DAPs regions from Rho-GFP/+ IS-enriched retina immunolabeled with NbGFP-A647 (magenta) and a CEP164 antibody (green). Centrin-2 antibody labeling marks the position of the CC (white), and residual NbGFP-A647 signal labels the OS; both are captured as widefield fluorescence images. White arrows indicate the regions that are magnified in the adjacent image. Scalebar values match adjacent panels when not labeled.

### Golgi and post-Golgi trafficking proteins are localized at the inner segment plasma membrane in mouse rods

We next tested the super-resolution localization of proteins that have previously been associated with the Rho secretory pathway in rod IS. Rab11a, a post-Golgi small GTPase, was shown to mediate Rho trafficking in post-Golgi vesicles in frog rods (Mazelova et al., 2009a; Wang et al., 2012) and was previously localized to the IS in mouse retinas in a semi-diffuse, puncta-like pattern (Grossman et al., 2011; Reish et al., 2014; Ying et al., 2016). Here, WT retina cryosections were immunolabeled with a specific Rab11a antibody (He et al., 2020), and Rab11a was localized with confocal imaging to the IS, in the ONL surrounding the nuclei, and in the OPL (Fig 9 A). With SIM, Rab11a was localized as bright puncta throughout the OS, IS and ONL, and in individual rod ISs, many Rab11a+ puncta were observed to be co-localized with the STX3+ IS plasma membrane (Fig. 9 B). After puncta counting, the rate of IS plasma membrane associated Rab11a+ puncta was 45.8% ± 14.3% (SD) per mouse rod IS (n=955 puncta, n= 32 rods); the rest of the Rab11a+ puncta were internal/cytoplasmic. In STORM reconstructions, Rab11a molecules were localized into molecule clusters within the IS, which were isolated for visualization using a Voronoi tessellation clustering algorithm (Fig. 9 C).

**Figure 9.**
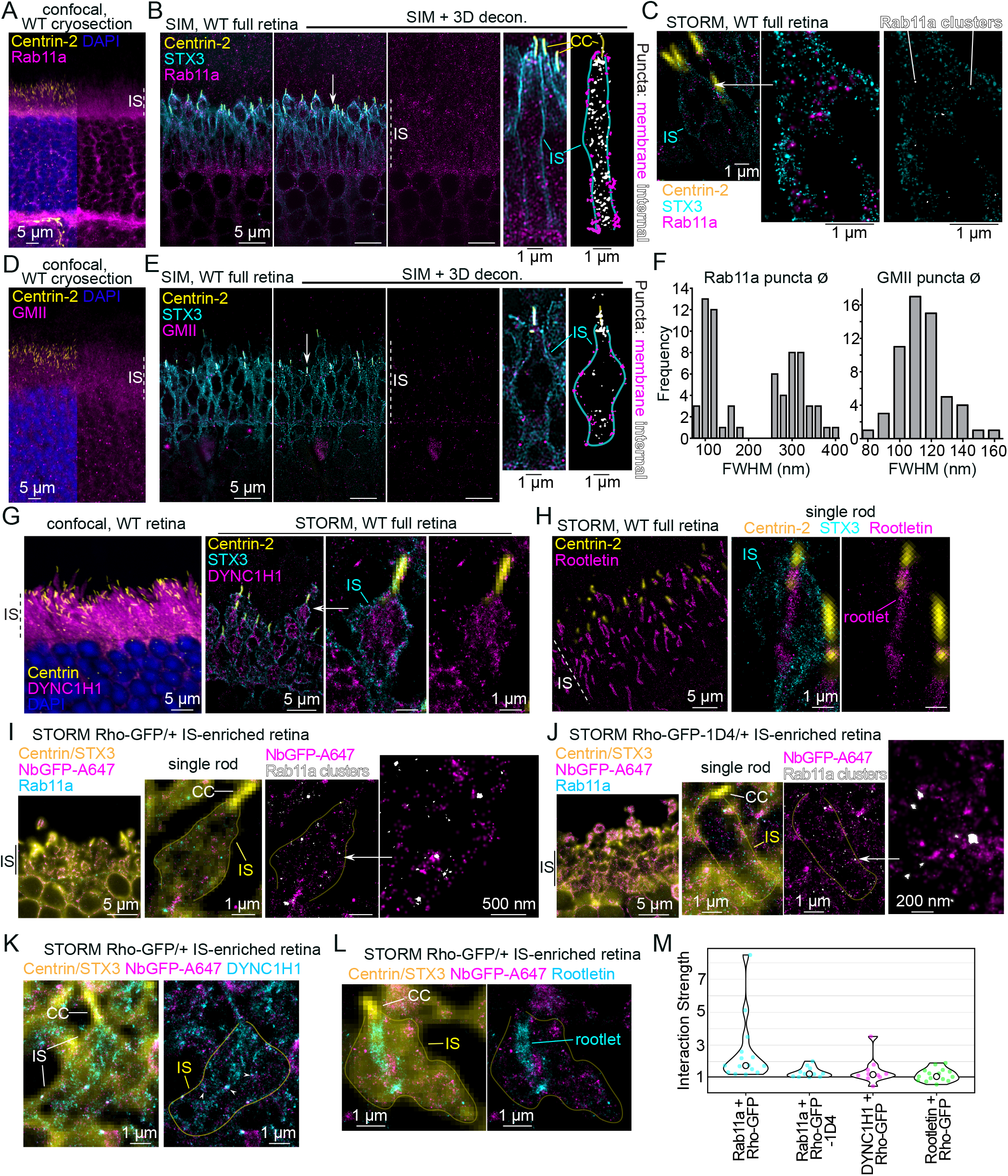
Rhodopsin colocalizes with Rab11a in mouse rod inner segments. (A) Z-projection of a WT retinal cryosection co-immunolabeled with centrin-2 and Rab11a antibodies and counterstained with DAPI. (B) SIM z-projection image of a WT retina co-immunolabeled with centrin-2, STX3 and Rab11a antibodies. The SIM retina image is also shown after 3D-deconvolution processing (SIM + 3D decon.) and Rab11a+ puncta are localized in the IS layer. A single rod cilium example is magnified, and a threshold image of the Rab11a channel is shown pseudocolored to depict puncta that are localized at the STX3+ IS membrane hull as magenta, and puncta that are localized internally or at the CC as white. (C) STORM reconstruction of a single rod from a WT retina immunolabeled as in (B). A magnified region is shown, and in the adjacent image, Rab11a+ clusters identified with Voronoi tessellation (see Methods) are in white. (D) Z-projection of a WT retinal cryosection co-immunolabeled with centrin-2 and GMII antibodies and counterstained with DAPI. (E) SIM images, including 3D decon. processed images, of a WT retina co-immunolabeled with centrin-2, STX3 and GMII antibodies. As in (B), a single rod example is shown, and in the adjacent image, membrane-localized GMII+ puncta are pseudocolored magenta, and internal (and ciliary) puncta are white. (F) Frequency plots for full width half maximum (FWHM) measurements of individual Rab11a and GMII IS-localized puncta from the SIM data represented in (B) and (E). For Rab11a FWHM values, n (number of puncta) = 67. For GMII FWHM values, n (number of puncta) = 58. (G) Confocal z-projection image of a WT retina cryosection immunolabeled with centrin and DYNC1H1 antibodies, as well as DAPI counterstaining. DYNC1H1+ fluorescence fills the IS layer. In the adjacent panel, a STORM reconstruction of a WT retina immunolabeled with centrin-2, STX3 and DYNC1H1 antibodies. A single rod example is shown. (H) STORM reconstruction of a WT retina immunolabeled with centrin-2, STX3 and rootletin antibodies. A single rod example is shown, and the ciliary rootlet is indicated. (I-M) STORM images of a (I) Rho-GFP/+ IS-enriched retina or (J) Rho-GFP-1D4/+ IS-enriched retina, each co-immunolabeled with NbGFP-A647, and centrin, STX3 and Rab11a antibodies. STORM reconstruction channels - NbGFP-A647 (magenta) and Rab11a (cyan) - are superimposed with the matching widefield fluorescence image of centrin/STX3 immunolabeling (combined, yellow). IS regions are indicated. A single rod example is shown, and the CC is indicated. In the adjacent image, Rab11a+ clusters identified with Voronoi tessellation are in white and the STX3+ IS hull is outlined in yellow. A white arrow indicates a further magnified region of Rho-GFP molecules localized around Rab11a clusters. Next, Rho-GFP was co-immunolabeled with (K) DYNC1H1 antibody or (L) Rootletin antibody. In both, the locations of the CC and the IS outline are indicated, and in (K) areas where Rho-GFP and DYNC1H1 molecules overlap are indicated with white arrowheads. In (L) the location of the ciliary rootlet is indicated. (M) STORM data from (I-L) conditions were used to perform Mosaic interaction analyses to test the colocalization between Rho-GFP molecules and the other immunolabeled target from the same rod IS. Interaction strength values are compared as violin plots (circles = median values and dashed lines = mean values). N-values, corresponding to the number of rods from each condition, are: Rab11a vs Rho-GFP, n=15; Rab11a vs Rho-GFP-1D4, n=11; DYNC1H1 vs Rho-GFP, n=10; Rootletin vs Rho-GFP, n=15. In all panels, white arrows indicate regions that are shown magnified. Scalebar values match adjacent panels when not labeled.

Next, although the GM130+ cis-Golgi was prominently localized in the myoid region of the IS (Fig. 5 B); the localization of Golgi alpha-mannosidase II (GMII), a *medial/trans*-Golgi glycoside hydrolase that was previously shown to process Rho protein in Golgi (Tian et al., 2014), was localized throughout the IS of mouse retinas using confocal microscopy (Fig. 9 D). In SIM images, GMII was also localized in discrete puncta within rod ISs, albeit less densely than Rab11a+ IS puncta (Fig. 9E); however, more GMII+ puncta were co-localized with the IS plasma membrane. The rate of IS membrane associated GMII+ puncta was 73.8% ± 19.2% (SD) (n= 303 puncta, n= 27 rods). Using SIM with 3D deconvolution, the diameters (⌀) of single, isolated IS Rab11a+ and GMII+ puncta were measured. Based on diameter, Rab11a+ puncta were distributed in 2 groups: <200 nm ⌀ puncta (mean ⌀ = 126.7 nm ± 23.5 nm, n = 34 puncta) and >200 nm ⌀ puncta (mean ⌀ = 320.8 nm ± 33.8 nm, n = 34 puncta), while GMII+ puncta were normally distributed as 1 group (mean ⌀ = 119.6 nm ± 14.5 nm, n = 58 puncta) (Fig. 9 F).

Together these results indicate that vesicular Golgi and post-Golgi trafficking organelles are targeted to the IS PM in mouse rods. For comparison, cytoplasmic dynein-1 and the ciliary rootlet were also localized in the mouse IS with super-resolution fluorescence. Cytoplasmic dynein-1 is essential in rods as the putative motor complex for intracellular cytoplasmic transport (Tai et al., 1999; Insinna et al., 2010; Dahl et al., 2021b; a). An antibody targeting the force-enerating heavy chain of the dynein-1 complex, DYNC1H1, was used to localize dynein-1 throughout the entire rod inner segment layer, and with STORM, DYNC1H1 molecules were localized in a homogenous distribution throughout the IS (Fig. 9 G). An antibody targeting rootletin, the core protein of the ciliary rootlet, was used to reconstruct the rootlet in rods with STORM (Fig. 9 H).

### Rhodopsin colocalizes with Rab11a in rod inner segments

STORM was used to test the co-localization of Rho-GFP with Rab11a, DYNC1H1 and Rootletin in IS-enriched Rho-GFP/+ retinas. In reconstructed mouse ISs, a fraction of Rho-GFP molecules were localized surrounding Rab11a+ molecule clusters (Fig. 9 I, Fig. S5 A). Similarly, in IS-enriched Rho-GFP-1D4/+ rods, STORM-reconstructed Rho-GFP-1D4 molecules partially accumulated around Rab11a+ clusters in the IS (Fig. 9 J, Fig. S5 B). Rho-GFP also partially co-localized with the more homogenously distributed DYNC1H1+ STORM molecules in the IS (Fig. 9 K, Fig. S5 C); however, Rho-GFP was not consistently co-localized with the ciliary rootlet (Fig. 9 L, Fig. S5 D). STORM colocalization from these mouse rod IS reconstructions was then quantified using the Mosaic spatial interaction analysis, and the interaction strength scores, normalized to random molecule distributions, were calculated (Fig. 9 M). From these aggregated scores, one sample t-tests were again performed to compare the normalized data to 1. Using this test, Rab11a + Rho-GFP and Rab11a + Rho-GFP-1D4 data were statistically greater than 1 (two-tailed P-values: Rab11a + Rho-GFP vs 1 P= 0.0089, Rab11a + Rho-GFP-1D4 vs 1 P= 0.0051) indicating a non-random co-localization between Rho and Rab11a in the IS. Although we observed partial co-localization between Rho-GFP and DYNC1H1 with STORM, the overall abundant and diffuse pattern of IS DYNC1H1 is likely why the interaction strength with Rho molecules was not distinguishably higher than random.

## DISCUSSION

In this study, we developed an optimized approach for localizing Rho in the IS of mouse rod photoreceptor cells. Rho is the most abundant protein in the membrane discs of rod OS cilia (Lyubarsky et al., 2004), which are constantly renewed through a robust network of protein translation and protein trafficking pathways in the IS. Based on our localization findings, including single molecule localization and spatial analysis data, we have established that Rho localized prominently at the IS plasma membrane in an even distribution along the entire boundary of the IS, including along the ellipsoid and myoid. IS plasma membrane Rho formed a string of localized fluorescence that was continuous with the CC plasma membrane. Rho co-localized at the IS plasma membrane with the t-SNARE protein STX3, and a significant fraction of Rab11a-positive and GMII-positive fluorescent puncta, as putative Golgi and post-Golgi vesicles, also localized at the IS hull. Combined with our co-localization of Rho with Rab11a, these findings suggest that after Golgi processing, new Rho proteins are trafficked within vesicles to the IS plasma membrane as a target/destination membrane. Figure 10 is a diagram of this model that also depicts Rho trafficking to the CC and then ultimately to the OS base within the IS plasma membrane through a lateral transport mechanism.

**Figure 10.**
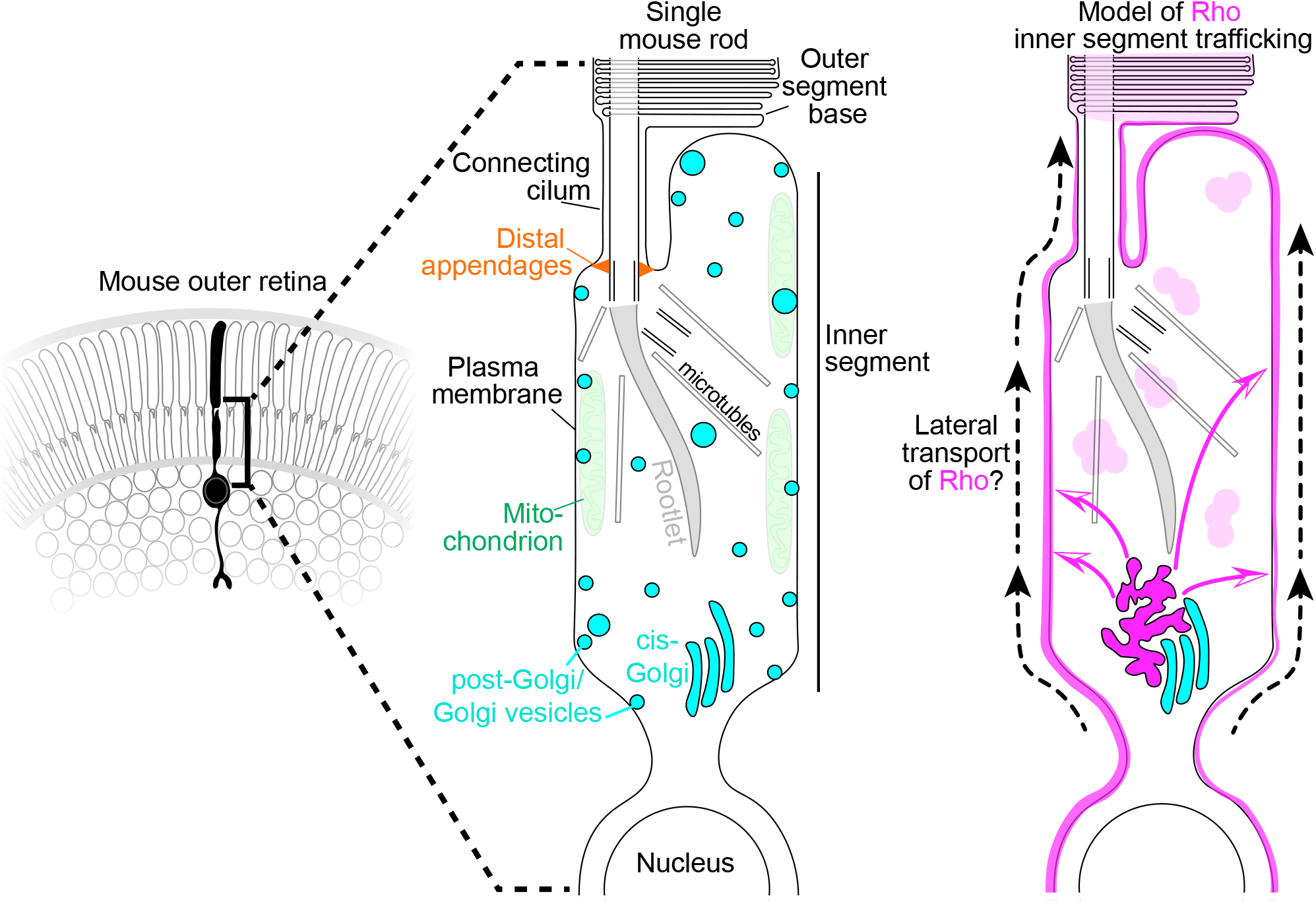
Diagram of the mouse rod inner segment architecture and the localization pattern of rhodopsin.

Such a lateral transport mechanism has been previously described for ciliary GPCRs in cultured mouse cells, in which Smoothened and the D1-type dopamine receptor proteins were shown to transport laterally between the plasma membrane and the primary cilia (Milenkovic et al., 2009; Leaf and Von Zastrow, 2015). Furthermore, dense rod IS plasma membrane localization of Rho was previously reported in pig retina (Röhlich et al., 1989), in mouse and cow rods (Jan and Revel, 1974), and in developing rat rods (prior to OS formation) (Nir et al., 1984). Further visualization of the Rho in IS in WT mammalian retina has likely been obscured in other reported imaging data by the overwhelming abundance of Rho in the OS and insufficient labeling of Rho in the IS. The literature does, however, contain many examples of Rho mislocalization to both the IS plasma membrane and the ONL in retinal degeneration studies using mammalian models. For example, the RP mutant Q344ter-Rho is mislocalized to the IS plasma membrane in transgenic mutant mice (Sung et al., 1994; Concepcion and Chen, 2010). A similar widespread IS Rho mislocalization was also seen in a RPGR mutant dog model of X-linked RP (Beltran et al., 2006), in a knockout mouse for the CC-localized and JBTS-associated Tmem138 membrane protein (Guo et al., 2022), and after experimental retinal detachment in cats (Fariss et al., 1997). In each example, a perturbation to OS trafficking likely generates a Rho localization imbalance between the OS vs IS, which then highlights the IS population of Rho.

Our model for IS Rho trafficking in mouse rods (depicted in Figure 10) deviates from the highly polarized vesicular trafficking pathway in frog rods, in which Rho-containing vesicles are directed to fuse to the IS plasma membrane directly adjacent to the CC at the ciliary base (reviewed in (Deretic et al., 2021)). Species differences between mouse and frog rods (Pearring et al., 2013; Ying et al., 2016), which likely contribute to IS trafficking differences, include photoreceptor cell size, CC length, IS mitochondria density, and the absence of ciliary rootlet in frogs. However, we have identified commonalities between the two rod trafficking pathways, including roles for Rab11a, STX3 and SNAP25, which are highlighted below.

SIM localization of both Rab11a+ and GMII+ fluorescent puncta to the IS plasma membrane (Fig. 9 A-F) provided further evidence that the mouse rod IS plasma membrane is an active site for protein trafficking. Based on diameter (⌀) Rab11a puncta were split into 2 groups, a group with a mean ⌀ ∼125 nm and another with a mean ⌀ ∼320 nm. GMII puncta were all in the same size range (mean ⌀ ∼120 nm). Interestingly, distal IS vesicles identified in rod cryo-EM tomograms ranged from 30-100 nm in diameter (Gilliam et al., 2012), while the majority of purified intracellular IS vesicles from frog rods were ∼300 nm in diameter (Deretic and Papermaster, 1991); however, because of our indirect fluorescent labeling for SIM, we cannot make direct size comparisons. Next, our STORM colocalization results with Rho and Rab11a (Fig. 9 I, J and Fig. S5 A, B) support a potential role for this Rab GTPase in IS Rho trafficking in mouse retina as previously described (Reish et al., 2014). Therefore, Rab11a+ fluorescent puncta may represent vesicles that are bound for SNARE-mediated vesicle fusion at the IS plasma membrane; however, they may alternatively mediate endocytotic events or be involved in a recently described exocytosis of ∼150 nm microvesicles from the IS plasma membrane (Lewis et al., 2022).

Any vesicular fusion events at the mouse rod IS plasma membrane would require STX3, which densely lines the IS plasma membrane of mouse rods (Fig. 4 B). Here we provide further demonstration of an essential interaction between Rho and STX3 using SIM and STORM colocalization and co-IP; an interaction which was previously established in rod-specific STX3 knockout mice (Kakakhel et al., 2020). In frog rods, STX3 and SNAP25 also line the IS plasma membrane (Mazelova et al., 2009b). Furthermore, the C-terminal SNARE domain of STX3 was identified as the IS retention sequence in frog rods that kept STX3 localized to the IS and prevented OS accumulation (Baker et al., 2008). Finally, while VAMP7 was identified as the v-SNARE in complex with STX3 and SNAP25 in frog rods (Kandachar et al., 2018), the functional v-SNARE in the mouse IS has not been identified (Zulliger et al., 2015).

We observed no consistent internal Rho localization in the ellipsoid IS or near the BB in mouse rods. Internal Rho that was sporadically localized in the ellipsoid IS may correspond to newly-synthesized Rho in the rough endoplasmic reticulum (ER). The ER has been previously localized in relatively close proximity to the BB in the distal mouse IS (Gilliam et al., 2012; Robichaux et al., 2022). In the myoid IS, on the other hand, we consistently observed bright puncta of Rho that did not co-localize with the GM130+ *cis*-Golgi puncta (Fig. 4 C). Instead the Rho fluorescence density was directly adjacent and possibly intercalated with the *cis*-Golgi. This finding suggests the existence of different *cis*-Golgi domains. Furthermore, the myoid rich Rho-Golgi complex in mouse rods may be analogous to Golgi exit sites in frog rods, which are also Golgi-adjacent accumulations of Rho protein in the IS (Deretic et al., 2005; Mazelova et al., 2009a).

Distal to the IS, we also observed CC membrane localized Rho using fluorescence (Fig. 3 B, Fig. 5 A), which had previously only been observed using post-embedding immuno-EM or cryo-immuno-EM (Liu et al., 1999; Tai et al., 1999; Wolfrum and Schmitt, 2000; Hagstrom et al., 2001; Burgoyne et al., 2015; Guo et al., 2022). We were also interested in visualizing Rho localization at the BB, and more specifically the DAPs, which is a critical boundary for any IS material to enter the CC. Our super-resolution localization of Rho surrounding the DAPs (Fig. 8) indicates that Rho is not integrated into the DAP structure and may only interact with CEP164 and other DAP proteins indirectly. Nonetheless, the DAPs are a structural barrier between the IS and CC (Wensel et al., 2021), and in a recent study, CEP164 conditional knockout mice had impaired OS disc formation defects likely due to decreased IFT protein recruitment to the BB and CC (Reed et al., 2022). Although the role of IFT transport in rods is currently unclear (Jiang et al., 2015), future efforts are needed to determine if plasma-membrane associated Rho that surrounds the DAPs is then coupled to the IFT complex along the CC.

Although we focused on the Rho localization in the IS of mouse rods, Rho was also located in the ONL (Fig. 2 E and Fig. 3 B), where it colocalized with STX3 (Fig. 4 C); however, because the rod nuclei are so prominent in this region, a suitable nuclear membrane marker will be needed for future ONL super-resolution fluorescent localization studies. Future studies will also be needed to connect our IS localization of Rho with other previously identified mouse IS trafficking regulators, including PrBP/δ (Zhang et al., 2007), IFT20 (Keady et al., 2011), myomegalin (Overlack et al., 2011), and spectrin βV (Papal et al., 2013). Nevertheless, the results presented here offer new insight into the organization of the mouse IS and may serve as the basis of a full mapping of IS trafficking processes using super-resolution fluorescence, live imaging and other complimentary techniques.

## MATERIALS AND METHODS

### Animals

All WT mice were C57BL/6 between the ages of 3 weeks and 6 months. The Rho-GFP and Rho-GFP-1D4 knock-in mice were previously described (Chan et al., 2004; Robichaux et al., 2022) and also C57BL/6. Mice were kept on a 12 hour light/dark cycle; however, mice used for SIM, STORM and all western blotting experiments were dark adapted the night before experiments and euthanized immediately after coming into the light the next morning to normalize the light exposure and timing of all retina samples. All co-IP conditions were repeated 3 times with retina samples from 3 different mice. All SIM and STORM conditions were repeated from multiple sections from at least 2 mice. Mice from both sexes were used indiscriminately All experimental procedures involving mice were approved by the Institutional Animal Care and Use Committee of West Virginia University.

### Antibodies and Labeling Reagents

The following primary antibodies were used in this study: anti-centrin (Millipore Cat# 04-1624, RRID:AB_10563501), anti-centrin-2 (BioLegend Cat# 698601, RRID:AB_2715793), anti-centrin-2 (Proteintech Cat# 15877-1-AP, RRID:AB_2077383), anti-STX3 (Millipore Cat# MAB2258, RRID:AB_1977423), anti-STX3 (Proteintech Cat# 15556-1-AP, RRID:AB_2198667), anti-Rho-C-1D4 (Millipore Cat# MAB5356, RRID:AB_2178961), anti-Rho-N-4D2 (Millipore Cat# MABN15, RRID:AB_10807045), anti-Rho-N-GTX (GeneTex Cat# GTX129910, RRID:AB_2886122), anti-ROM1 (Proteintech Cat# 21984-1-AP, RRID:AB_2878961), anti-Rab11a (Abcam Cat# ab128913, RRID:AB_11140633), anti-Tubulin/TUBB5B (Sigma-Aldrich Cat# T7816, RRID:AB_261770), anti-phosducin (custom-made rabbit polyclonal antibody, gift from Dr. Maxim Sokolov), anti-SNAP25 (BioLegend Cat# 836304, RRID:AB_2566521), anti-GM130 (BD Biosciences Cat# 610822, RRID:AB_398141), anti-mannosidase II/GMII (Abcam Cat# ab12277, RRID:AB_2139551), anti-DYNC1H1 (Proteintech, 12345-1-AP), anti-Rootletin (Millipore Cat# ABN1686, RRID:AB_2893142), and anti-ROS-GC1 (Santa Cruz Biotechnology Cat# sc-376217, RRID:AB_10991113).

The following secondary antibodies were used in this study: F(ab’)2-goat anti-mouse IgG Alexa 647 (Thermo Fisher Scientific Cat# A48289TR, RRID:AB_2896356), F(ab’)2-goat anti-rabbit IgG Alexa 647 (Thermo Fisher Scientific Cat# A-21246, RRID:AB_2535814), F(ab’)2-goat anti-mouse IgG Alexa 488 (Thermo Fisher Scientific Cat# A-11017, RRID:AB_2534084), AffiniPure F(ab’)2-goat anti-rat IgG (H+L) Alexa 488 (Jackson ImmunoResearch Labs Cat# 112-546-003, RRID:AB_2338364), F(ab’)2-goat anti-mouse IgG CF568 (Biotium Cat# 20109, RRID:AB_10557119), F(ab’)2-goat anti-rabbit IgG CF568 (Biotium Cat# 20099, RRID:AB_10563029), F(ab’)2-goat anti-mouse IgG Alexa 555 (Thermo Fisher Scientific Cat# A-21425, RRID:AB_2535846), F(ab’)2-goat anti-rabbit IgG Alexa 555 (Thermo Fisher Scientific Cat# A-21430, RRID:AB_2535851), anti-mouse IgG sdAb – FluoTag-X2 – Alexa 647 (Nanotag, #N2002-AF647-S), Nanogold-Fab’ goat anti-mouse (Nanoprobes Cat# 2002, RRID:AB_2637031), and Nanogold-Fab’ goat anti-rabbit (Nanoprobes Cat# 2004, RRID:AB_2631182).

To generate the GFP nanobody Alexa 647 conjugate, NbGFP-A647, the pGEX6P1-GFP plasmid (RRID:Addgene_61838) was transformed into BL21 (DE3) competent cells (Thermo Scientific) to grow recombinant GST-nanobody in Luria Broth (LB) supplemented with 50 µg/ml carbenicillin. At mid-log phase 1000 µM isopropyl β–d-1-thiogalactopyranoside (IPTG) was added to induce protein expression at 16 C. Bacterial pellets were frozen at -80°C, thawed, and incubated in the following detergent buffer: 2% Triton X-100, 0.2 mg/ml lysozyme, 20 U/ml Pierce Universal Nuclease (Thermo Scientific), 50 mM NaCl, 1 mM MgCl2, 10 mM beta-mercaptoethanol (BME), 50 mM Tris (pH 8.0), EDTA-free protease inhibitor cocktail (Bimake.com) for 0.5-1 h on ice. Before centrifugation, 0.5% deoxycholate and 250 mM NaCl was added to the lysates, which were then centrifuged for 30 min at 12,000 xG at 4°C in a fixed-angle rotor. The cleared supernatant was loaded onto a GE ÄKTA start system fast protein liquid chromatography (FPLC) system using 2x 1 ml GSTrap columns (Cytiva #17528105) with a flow rate of 0.5 ml/min. 20 mM glutathione in 50 mM TRIS/HCl, pH 8.0, was used to elute the GST-nanobody. Eluates nanobody protein samples were buffer-exchanged into proteolysis buffer (25 mM HEPES, 150 mM NaCl, 1 mM DTT, 1 mM EDTA, pH 7.5) and incubated with 3 mg of Biotin HRV-3C protease (Sigma, cat# 95056) to remove the GST-tag, and then FPLC and the same GSTrap columns was used again to remove the cleaved tags. Purified nanobody was buffer-exchanged into PBS, pH 8.5, validated with SDS-PAGE and Coomassie blue staining, and quantified by UV spectrophotometry.

Purified GFP nanobody was incubated with Alexa Fluor 647 NHS Ester (Thermo Fisher Scientific Cat# A20006) at a molar 1:4 ratio, and the mixture was incubated for 2 h at RT, protected from light, with occasional tapping. During this time, 2x 5 ml PD-10 G-25 desalting columns (Cytiva Cat# 17085101) were equilibrated with 1x PBS (pH 8.5). After incubation, the conjugate mix was added to one column and 200 µl 1x PBS (pH 8.5) fractions were collected. A NanoDrop 2000 (Thermo Scientific) was used to screen the absorbance profiles at 280 nm and 647 nm to identify the nanobody-Alexa 647 fractions. Those positive fractions were purified on the second equilibrated G-25 desalt column and screened for absorbance. The nanobody-Alexa 647 positive fractions from the second desalting purification were combined, stored at 4°C, and used as the NbGFP-A647 reagent for immunolabeling.

### Retinal immunofluorescence

For immunofluorescence and confocal microscopy, mouse eyes were enucleated and a small puncture was made in the cornea and eyes was subsequently fixed for 15 min in 4% PFA at RT. The cornea and lens were then completely removed to form an eye cups, which were further fixed in 4% PFA for 45 min at RT. Following incubation in 30% sucrose for cryopreservation, eye cups were transferred to a 1:1 mix of 30% sucrose and optical cutting temperature medium (OCT) for additional cryopreservation. Eye cups were frozen in cryomolds and 16 µm sections were collected on either a Leica Cryostat CM1850 or a Medical Equipment Source 1000+ Cryostat and collected on Superfrost slides (VWR, Cat# 48311-703). For immunolabeling, sections were quenched with 100 mM glycine diluted in 1x PBS for 10 min at RT. Sections were then incubated with blocking solution: 10% normal goat serum (NGS) (Fitzgerald), 0.3% Triton X-100, 0.02% sodium azide, diluted in 1x PBS, for 1 h at RT. 1 µg – 5 µg of primary antibodies were diluted in 200 µl the same block solution and added to the sections for overnight probing at 4°C in a humidified chamber. The next day, sections were washed and probed with 1 µg of fluorescent secondary antibodies diluted 200 µl of the same blocking solution for 1.5 h at RT. After washing, sections were counterstained with 0.2 µg/ml 4′,6-diamidino-2-phenylindole (DAPI) (Thermo Fisher Cat# 62248) diluted in 1xPBS for 15-30 min at RT. Sections were post-fixed in 4% PFA for 5 min and mounted with ProLong Glass Antifade Mountant (Thermo Fisher Scientific Cat# P36980). Confocal scanning was performed at RT on either a Nikon C2 confocal microscope equipped with photomultiplier detectors using 40x Plan Fluor, NA 1.3 and 100x Plan Apo, NA 1.45 oil immersion objectives, or a Nikon A1R confocal microscope equipped with GaAsP and photomultiplier detectors using a 40x Plan Fluor, NA 1.3 objective. Confocal images were acquired using NIS-Elements software and processed using Fiji/ImageJ (Schindelin et al., 2009).

For SIM and STORM immunofluorescence, dissected mouse retinas were dissected in ice cold buffered Ames media (Sigma, Cat# A1420) and either lightly fixed in 4% PFA diluted in Ames’ for 5 min on ice (for whole retina samples), or immediately underwent retinal peeling for IS enrichment. The peeling procedure is adapted from (Rose et al., 2017, 2021). Dissected retinas for peeling were first bisected and then trimmed into rectangular retinal slices. 5.5 cm filter paper (VWR, Cat# 28310-015) was cut into small squares, a piece of filter paper was added to dish of Ames’ media containing the retinal slices, and the slices was oriented with the OS facing the filter paper. The filter paper with retina slice attached were carefully lifted out of solution and blotted dry filter side down on a paper towel before being transferred back into the Ames’ media, and this blotting procedure was repeated 4 times. After blotting, the retina was carefully peeled away from the filter paper using forceps. This process constituted a single peel. In total, the retina slices were peeled 8 times (for IS-enriched retinas) before immediately being lightly fixed in 4% PFA diluted in Ames’ for 5 min on ice. After fixation, retina samples (either whole retinas or IS-enriched retinas) were quenched in 100 mM glycine at 4°C and incubated in 1 mL SUPER block solution: 15% NGS, 5% bovine serum albumin (BSA) (Sigma, Cat# B6917) + 0.5% BSA-c (Aurion, VWR, Cat# 25557) + 2% fish skin gelatin (Sigma, Cat# G7041) + 0.05% saponin (Thermo Fisher, Cat# A1882022) + 1x protease inhibitor cocktail (GenDepot, Cat# P3100-005), in half dram vials (Electron Microscopy Sciences, Cat# 72630-05) or low-adhesion microcentrifuge tubes (VWR, Cat# 49003-230) for 3 h at 4°C. 1 µg – 5 µg of primary antibodies or 6 µg NbGFP-A647 was added to 1 ml of the blocking solution for probing at 4°C for 24 h with mild agitation. The next day, a second dose of Rho antibodies or NbGFP-A647 (same amounts as the day prior) was added to improve retinal labeling penetration, and the retinas were probed at 4°C for an additional 48 h with mild agitation (for 72 h total of primary immunolabeling). Retinas were washed 6 times for 10 min each in 2% NGS diluted in Ames’ prior to probing with 4 µg – 8 µg of secondary antibodies diluted in 1 ml of 2% NGS in Ames’ + 1x protease inhibitor cocktail at 4°C for 12-16 h with mild agitation. Retinas were then washed 6 times, 5 minutes each in 2% NGS diluted in Ames’ and fixed in 2% PFA + 0.5% glutaraldehyde diluted in 1xPBS for 30 min at 4°C with mild agitation. Retinas were then dehydrated in an ethanol series with the following steps of pure ethanol diluted in water: 50%, 70%, 90%, 100%, 100%; each step for 15 minutes in half dram vials on a room temperature roller. Dehydrated retinas were then embedding in Ultra Bed Low Viscosity Epoxy resin (EMS Cat# 14310) in the following series of steps: 1:3 resin to 100% ethanol for 2 h rolling at RT; 1:1 resin to 100% ethanol for 2 h rolling at RT; 3:1 resin to 100% overnight (∼16 h) rolling at RT; two steps of full resin (no ethanol), 2 h each, rolling at RT. Resin-embedded retina samples were mounted in molds and cured in baking oven set to 65°C for 24 h. A Leica UCT ultramicrotome and glass knives were used to make 0.5 µm – 1 µm thin retinal cross sections.

For SIM, thin retina resin sections were collected onto #1.5 coverslips (VWR, Cat# 1152222), which were then mounted onto a plain glass slides with ProLong Glass and cured at RT for at least 2 full days protected from light. For STORM, thin retina resin sections were collected in 35 mm glass-bottom dishes (MatTek Life Sciences, Cat# P35G-1.5-14-C), chemically etched in a mild sodium ethoxide solution (∼1% diluted in pure ethanol for 0.5 – 1.5 h) as previously described (Robichaux et al., 2019). Prior to STORM imaging, etched sections were mounted in a STORM imaging buffer adapted from (Albrecht et al., 2022): 50 mM Tris (pH 8.0), 10 mM NaCl, 10 mM sodium sulfite, 10% glucose, 40 mM cysteamine hydrochloride (MEA, Chem Impex/VWR, Cat# 102574-806), 143 mM BME, and 1 mM cyclooctatetraene (Sigma Cat# 138924), under a #1.5 glass coverslip that was sealed with quick-set epoxy resin (Devcon).

An alternative protocol was used for SIM imaging of GFP fluorescence in Rho-GFP/+ and Rho-GFP-1D4/+ retinal cryosections. Enucleated eyes were immersion fixed in 4% PFA diluted in 1x PBS for 1h at RT and incubated in a hydrogel solution: 4% acrylamide (Sigma, Cat# A4058), 2% bis-acrylamide (Alfa Aesar Cat# J63265), 2.5% VA-004 (TCI America), overnight at 4°C. The hydrogel was polymerized for 5 min at 37°C and degassed for 15 min at RT. Hydrogels were cryoprotected with sucrose, embedded in OCT and frozen. 2 µm cryosections were collected on Superfrost slides, rinsed in 1xPBS, mounted in ProLong Glass Antifade Mountant, and cured for 2 days at RT before SIM.

### SIM

A Nikon N-SIM E microscope system equipped with a Hamamatsu Orca-Flash 4.0 camera was used for imaging at RT with a SR HP Apochromat TIRF 100X, NA 1.49 oil immersion objective. NIS-Elements Ar software was used for image acquisition, and Z-projections were collected from regions of interest (containing 5-15 slices), using a 0.2 µm Z-section thickness. Samples were imaged using 488 nm, 561 nm, and 647 nm lasers with 15 grating pattern images. SIM reconstructions were performed in NIS-Elements software and assessed for quality based on the fast Fourier transform (FFT) images from the reconstructions. 3D deconvolution was also performed in Nikon NIS-Elements software using Automatic deconvolution mode.

### STORM

All STORM acquisitions were performed at RT on a Nikon N-STORM 5.0 system equipped with an Andor iXON Ultra DU-897U ENCCD camera with a SR HP Apochromat TIRF (total internal reflection fluorescence) 100X, NA 1.49 oil immersion objective. The system also features a piezo Z stage,100 mW 405 nm, 488 nm and 561 nm laser lines and a 125 mW 647 nm laser line, as well as a Lumencor SOLA Light Engine epifluorescence light source. For STORM acquisition, Nikon NIS-Elements software was used, and sections and imaging fields of interest were located using widefield epifluorescence. All STORM was performed in 2D mode using 40 µm x 40 µm field of view dimensions. 561 nm and 647 nm lasers were used at low laser power to identify spots with multiple bright CC, and then pre-STORM images were acquired using epifluorescence and lower laser power. GFP and Alexa 488 fluorescence were photobleached using 488 nm laser at maximum power prior to using the 561 nm and 647 nm lasers at 90%-100% max power to initiate CF568 and Alexa 647 photoswitching. 2x laser magnification was used for optimal photoswitching, and all STORM acquisitions was performed near the TIRF critical angle; the TIRF angle and direction values were adjusted for each sample to optimize signal-to-noise in each imaging field. STORM data were acquired at ∼33 frames per second, and 40,000 frames were collected per channel for each STORM acquisition. 561 nm and 647 nm frames were collected sequentially. STORM analysis was performed using Nikon NIS-Elements software with analysis parameters matching the “single molecule” settings from (Robichaux et al., 2019) to detect only bright, individual photoswitching events, as follows: Minimum PSF height: 1,000, Maximum PSF height: 65,000, Minimum PSF Width: 200 nm, Maximum point spread function (PSF) Width: 400 nm, Initial Fit Width: 300 nm, Max Axial Ratio: 1.15, Max Displacement: 1 pixel. The resulting STORM reconstructions were populated as single molecule events with localization errors <20 nm. Chromatic aberration between channels was corrected using an X-Y bead warp calibration and drift correction was performed using an auto-correlation algorithm

### TEM

IS-enriched or full (unpeeled control) retinas for TEM were immediately fixed in 2% PFA + 2% glutaraldehyde + 4.5 mM CaCl_2_ in 50 mM MOPS buffer (pH 7.4) for 4-5 h at 4°C on a roller. Retinas were then incubated in half dram vials containing 1% tannic acid (EMS, Cat# 21700) + 0.5% saponin diluted in 0.1M HEPES (pH 7.5) for 1 h at RT, then rinsed 4 times with water before incubation with 1% uranyl acetate (EMS, Cat# 22400) diluted in 0.1 M maleate buffer (pH 6.0) for 1 h at RT. Retinas were then rinsed 4 times in water and then dehydrated in an ethanol series with the following steps of pure ethanol diluted in water: 50%, 70%, 90%, 100%, 100%; each step was for 15 min in half dram vials on a RT roller. Dehydrated retinas were then embedded in Ultra Bed Low resin and cured in resin molds as described for SIM/STORM retinas above.

70 nm ultramicrotome sections were cut using a Diatome Ultra 45° diamond knife and collected onto square 100 mesh copper grids (EMS, Cat# G100-Cu). Grids were post-stained in 1.2% uranyl acetate diluted in water for 4 min, rinsed 6 times in water, and allowed to completely dry before staining with either a Sato’s Lead solution (1% lead acetate (EMS, Cat# 17600), 1% lead citrate (EMS, Cat# 17800), 1% lead nitrate (EMS, Cat# 17900), 2% sodium citrate diluted (EMS, Cat# 21140) in water) or a lead citrate solution (EMS, #22410) for 4 min. Grids were then rinsed in water and dried overnight. Grids were imaged using a JEOL JEM 1010 transmission electron microscope equipped with a side-mounted Hamamatsu Orca-HR digital camera at 25,000x magnification.

### Immuno-EM

Retinas were dissected, IS-enriched and immunolabeled as described in the SIM/STORM section except the pre-fixation solution was 4% PFA + 2.5% glutaraldehyde in Ames’, nanogold-conjugated secondaries were used (5 - 7.5 µg), and the post-fixation solution was 2% PFA + 2% glutaraldehyde + 4.5 mM CaCl_2_ in 50 mM MOPS buffer (pH 7.4). Retinas were then rinsed with water and enhanced using HQ Silver Kit (Nanoprobes, Cat# 2012) reagents in half dram vials for 4 min at RT with agitation. Enhanced retinas were then immediately rinsed with water, incubated in 1% tannic acid + 0.5% glutaraldehyde in 0.1 M HEPES (pH 7.5) for 1 h on a RT roller, rinsed with water, incubated 1% uranyl acetate in 0.1 M maleate buffer (pH 6.0) for 1 h on a room temperature roller, and rinsed a final time with water. Retinas were then ethanol dehydrated and resin embedded in Ultra Bed resin as outlined in the TEM section. 100 nm ultramicrotome sections were cut using a Diatome Ultra 45° diamond knife and collected onto square 100 mesh copper grids. Grids were imaged as for TEM.

### Image processing and spatial analysis

Fiji/ImageJ was used to generate confocal and SIM z-stack projections, adjust whole image brightness and contrast adjustments for clarity, and generate row average intensity profiles (using the “Plot Profile” function). For SIM fluorescent puncta analyses, puncta membrane localization was scored using SIM z-projection images. Puncta diameters were acquired by boxing single puncta in Fiji/ImageJ, acquiring row average intensity profiles and using the “Add Fit…” function to fit Gaussian functions to the profiles, which were exported to a custom code in Mathematica v13.1 (Wolfram) to determine full width half maximum (FWHM) diameter values.

STORM reconstruction data were processed in NIS Elements Ar v5.30.05 (Nikon) using the N-STORM Analysis modules. Rod IS hulls were manually drawn on STORM reconstructions using the “Draw Polygonal ROI…” tool, and molecule coordinates within IS hulls were exported for further processing. Widefield images were superimposed onto STORM reconstructions and exported as image files to be adjusted, scaled and cropped for visualization in Fiji/ImageJ. STORM spatial analysis was performed using custom code in Mathematica v13.1.0.0 (Wolfram). Molecule coordinates corresponding to single rod IS hulls were imported and plotted using the ListPlot function, and the IS hull was defined using the ConcaveHullMesh function on the STX3 coordinates. Random points were generated using the RandomReal function, with points outside the IS hull discarded. Nearest distance to hull measurements were calculated using the SignedRegionDistance function and graphed as frequency plots or CDF plots using the PDF or CDF functions, respectively. Finally, ≤0.1 µm, ≤0.2 µm, or ≤0.3 µm frequency values were calculated as the number of molecules with nearest distance to hull within the indicated range, divided by the total number of molecules.

The Mosaic IA Fiji/ImageJ plugin (Shivanandan et al., 2013) was used to perform the STORM spatial interaction analysis to quantify STORM colocalization. Coordinates for each channel from an IS hull were uploaded (with the STX3 channel serving as the reference channel). The following settings were used: Grid spacing = 0.03, Kernel wt(q) = 0.001, Kernel wt(p) = based on the estimate provide by the program. The parameterized “Hernquist potential” was used to calculate the interaction strength values. Rab11a clusters were visualized in STORM reconstructions in NIS-Elements Ar using as Voronoi tessellations using the Molecule Analysis feature within the N-STORM Analysis program; Voronoi parameters were set to cluster 3 or more molecules in a 10 nm maximum distance. Tessellations were visualized using the Render Clusters function and superimposed onto STORM reconstruction images in NIS-Elements.

### Western blotting

For retinal lysate western blotting, mouse retinas were dissected and residual retinal pigment epithelium and ciliary body material were removed. Single retinas were then frozen individually on dry ice for at least ten minutes before adding 200 µl of urea sample buffer: 6M urea, 140 mM SDS, ∼0.03% bromophenol blue diluted in 0.125 M Tris (pH 6.8) and supplemented with 360 mM BME. Samples were sonicated to lyse the retinas. Lysates were centrifuged 1 – 5 min and supernatants were collected. Samples were loaded onto Novex WedgeWell 10-20% Tris-Glycine, 0.1 mm, MiniProtein Gels (Invitrogen, Cat# XP10202BOX) along with the Precision Plus Dual Color ladder (Bio-Rad, Cat# 1610374) in Tris-Glycine-SDS running buffer (Bio-Rad, Cat# 1610772) for SDS-PAGE. Gels was transferred onto Immobilon-FL Transfer Membrane PVDF (pore size: 0.45 µm) (LI-COR Cat# 92760001) in Tris-Glycine Transfer Buffer (Bio-Rad Cat# 1610771) + 10% methanol. Membranes were subsequently blocked using Intercept Blocking Buffer (LI-COR, Cat# 927-6000) for 1 h and washed before primary antibody staining in 1xPBS + 0.1 Tween-20 (PBS-T) (antibodies were used at 1:500 - 1:5000 dilutions) for 1 h. Membranes were washed in PBS-T 3 times, 5 min each before secondary staining with Alexa 647 secondary antibodies (1:50,000 each) diluted in PBS-T for 1 h. Secondary antibodies used were F(ab’)2-goat anti-mouse IgG Alexa 647 and F(ab’)2-goat anti-rabbit IgG Alexa 647. Membranes were washed then imaged for fluorescence on an Amersham Typhoon scanner (GE) using both Cy5 and IR-Short fluorescence filter sets; the IR-short channel was optimal for imaging the protein ladder

### Co-immunoprecipitation

Retinas were dissected and IS-enriched as described above before each individual retina was frozen on dry-ice. 200 µl T-PER lysis buffer (Thermo Fisher Scientific, Cat# 78510) was added to frozen retinas prior to sonication. Retinal lysates were centrifuged and soluble supernatants were collected. 4 µl supernatant was saved as “input” fractions for SDS-PAGE (corresponding to 2% of a lysed retina). 20 µl of either anti-GFP Trap magnetic beads (Chromotek Inc. Cat# gtma) or 1D4-Ab-conjugated sepharose beads (∼30% slurry, gift from Dr. Theodore Wensel) was added directly to remaining lysates and incubated overnight at 4°C with agitation. Following overnight incubation, samples were pelleted and 4 µl supernatant was removed as the “unbound” fraction for SDS-PAGE (also corresponding to 2% of a lysed retina). For magnetic beads, a magnetic stand was used to pellet beads for washes. For agarose beads, samples were centrifuged to pellet the beads. All beads were washed a total of 4 times for 3 min each in PBS-T at RT. After washing, the “bound” sample was eluted from beads using 25 µl urea sample buffer supplemented with 360 mM BME. Half of the bound eluate sample was loaded alongside the corresponding input and unbound samples for SDS-PAGE in Novex WedgeWell 10 to 20% Tris-Glycine, 0.1 mm, MiniProtein Gels (Invitrogen) with Tris-Glycine-SDS running buffer (BioRad). Western blot transfer, immunolabeling and scanning steps were performed as described in the western blotting section.

### Deglycosylation assay

Full mouse retinas were dissected as described above. After freezing, 200 µl of RIPA lysis buffer (Alfa Aesar, Cat# J63306) supplemented with 1x protease inhibitor cocktail (GenDepot, Cat# P3100-005) was added directly to a frozen retina, which was sonicated and centrifuged as described above. Supernatant concentrations were measured using a NanoDrop 2000 (Thermo Fisher Scientific). 100 µg of lysates were deglycosylated using Protein Deglycosylation Mix II (NEB Cat# P6044S). In short, lysates were incubated with 1X Deglycosylation Buffer 2 for 10 min at 37°C, then 1X Deglycosylation Mix II (containing PNGase F) was subsequently added, and samples were incubated for 1 h at 37°C. The reaction was terminated by cooling lysates on ice for 5-10 minutes. Lysates were then further diluted to 50 µg in urea sample buffer supplemented with BME for western blotting.

### Statistical analysis

Mann-Whitney U tests were performed in Microsoft Excel by ranking the the values from each condition and calculating the U test and z test statistics (to derive p-values via hypothesis testing). One sample t-tests for comparisons to the hypothetical mean value of 1 were performed using GraphPad QuickCalcs (https://www.graphpad.com/quickcalcs/). Data were visualized for comparison as violin plots using PlotsOfData (Postma and Goedhart, 2019).

## SUPPLEMENTAL MATERIAL

Fig. S1 shows data testing the specificity of the NbGFP-A647 reagent and the deglycosylation assay western blots. Fig. S2 contains additional TEM images of IS-enriched retinas, SIM images of Rho localization in IS-enriched Rho-GFP-1D4/+ mouse retina sections, and additional SIM images of endogenous mouse Rho localization with the Rho-C-1D4 and Rho-N-4D2 antibodies. Fig. S3 is a supplemental figure for Fig. 6 showing additional STORM spatial analysis example data in addition to STORM data from IS-enriched Rho-GFP-1D4/+ mouse retina sections. Fig. S4 shows the normalized frequency <0.1 µm and <0.3 µm violin plot graphs for the STORM spatial analysis data and also the immuno-EM data. Fig. S5 is a supplemental figure for Fig. 9 compiling more colocalization example STORM images.

## DATA AVAILABILITY STATEMENT

All confocal microscopy image data are available in the published article. SIM raw data and STORM molecule list data are openly available via Mendeley Data at DOI: 10.17632/pv3ty3z68f.1, DOI: 10.17632/r4246hbxsw.1, and DOI: 10.17632/hmztjrwhtc.1. Original western blot scans are available in source data files.

## ACKNOWLEDGEMENTS

The authors thank Dr. Paolo Fagone and WVU Department of Biochemistry and Molecular Medicine Viral Core Facility for assistance with GFP nanobody purification. The authors thank Drs. Abigail Moye, Ezequiel Salido, Maxim Sokolov, and Theodore Wensel, as well as Robichaux lab members, for discussion and feedback during manuscript preparation. The authors declare no competing financial interests.

This work was supported by National Institute of Health research grant P20-GM144230 (M.A.R.) and a grant from the Knights Templar Eye Foundation (M.A.R.). Confocal and SIM imaging experiments were performed in the West Virginia University Microscope Imaging Facility, which has been supported by the WVU Cancer Institute and National Institute of Health research grants P20RR016440 and P30RR032138/P30GM103488. The Nikon A1R-SIM is supported with funding from U54GM104942 & P20GM103434.

## AUTHOR CONTRIBUTIONS

Conceptualization: K.N.H., S.C.E., M.A.R.; Investigation: K.N.H., S.C.E., L.A.S., E.F., L.M.R., M.A.R.; Resources: M.A.R.; Writing – original draft: M.A.R.; Writing – reviewing & editing: K.N.H., S.C.E., M.A.A., M.A.R., Visualization: M.A.A., M.A.R.; Software: M.A.A.; Project Administration: M.A.R.; Supervision: M.A.R. Funding Acquisition: M.A.R.

## ABBREVIATIONS

BB: basal body
BME: beta-mercaptoethanol
BSA: bovine serum albumin
co-IP: coimmunoprecipitation
CC: connecting cilium
DAPI: 4′,6-diamidino-2-phenylindole
EMS: Electron Microscopy Sciences
ER: endoplasmic reticulum
FWHM: full width half maximum
GPCR: G protein coupled receptor
IFT: intraflagellar transport
immuno-EM: immunoelectron microscopy
IS: inner segment
OCT: optimal cutting temperature medium
OS: outer segment
NbGFP-A647: GFP nanobody conjugated to Alexa 647
NGS: normal goat serum
PBS-T: 1xPBS + 0.1% Tween-20
PDC: phosducin
PSF: point spread function
RP: retinitis pigmentosa
Rho: rhodopsin
SIM: structured illumination microscopy
SNARE: soluble N-ethylmaleimide-sensitive factor attachment protein receptor
STORM: stochastic optical reconstruction microscopy
STX3: syntaxin 3
TEM: transmission electron microscopy

## FIGURE LEGENDS

**Figure S1.** (A, B) Western blot test of NbGFP-A647 immunolabeling specificity. (A) WT and Rho-GFP/+ retinal lysates used as either 0.5% or 2% total volume of 1 mouse retina. In Rho-GFP/+ lysates, a prominent and specific NbGFP+ band was found ∼60 kDa corresponding to monomeric Rho-GFP protein. Both WT and Rho-GFP/+ lysates contained Rho-C-1D4-positive bands corresponding to endogenous mouse Rho protein. Mouse Rho protein levels are apparently lower in Rho-GFP/+ lysates – corresponding to the one WT *Rho* allele in these mice. In addition, Rho-C-1D4 antibody does not immunolabel Rho-GFP protein from Rho-GFP/+ lysates. IB=immunoblot condition. (B) WT and Rho-GFP-1D4/+ retinal lysates used as 2% total volume of 1 mouse retina. In Rho-GFP-1D4/+ lysates specific NbGFP-A647+ and Rho-C-1D4+ bands are found ∼60 kDa corresponding to monomeric Rho-GFP-1D4 protein. As is (A), endogenous mouse Rho protein levels are lower in Rho-GFP-1D4/+ lysates. For all western blots, molecular weight marker sizes are indicated in kDa. (C) To test NbGFP-A647 immunolabeling specificity with immunofluorescence, 10 µm eye cup cryosections from Rho-GFP/+ mice (age P71) and from WT mice (age P60) were immunolabeled with NbGFP-A647 and counterstained with DAPI. NbGFP-647 specifically labels the outer segments (OS) in the Rho-GFP/+ section (magenta) and overlaps with GFP (green). No detectable NbGFP-A647 fluorescence was detected in the WT sections. Scalebar values match adjacent panels when not given. (D) Rho-GFP/+ or (E) Rho-GFP-1D4/+ adult retinal lysates treated with either Protein Deglycosylation Mix II from NEB (containing PNGase F) or buffer only. After treatment, 5 µg of total protein was loaded for SDS-PAGE, transfer and primary antibody labeling. A shift to lower molecular weight was observed for all Rho bands, including Rho-GFP bands from Rho-GFP/+ treated lysates probed with NbGFP-A647. In (D) Rho-GFP was also detected with Rho-N-4D2 immunolabeling; monomeric Rho-GFP bands are indicated with green arrows on this blot. In (E) Rho-GFP-1D4 was detected with Rho-C-1D4 immunolabeling and monomeric Rho-GFP-1D4 bands are indicated with green arrows.

**Figure S2.** (A) TEM images of Rho-GFP/+ retina slices that were either unpeeled, peeled 4 times or peeled 8 times (the IS-enriched condition). Images were pseudocolored to point out key rod structures as follows: OS = yellow, IS = magenta, connecting cilia (CC)/basal bodies = blue. Scalebar values match adjacent panels when not labeled. (B) Alternate TEM single rod examples from Rho-GFP/+ IS-enriched retinas. The IS plasma membrane is annotated with magenta arrows. (C) SIM images from IS-enriched Rho-GFP/+ retinas immunolabeled for NbGFP-A647 (magenta), STX3 (cyan), and centrin (yellow). To demonstrate Rho-GFP colocalization with STX3 at the IS plasma membrane, row average intensity plots are shown for portions of the IS from 2 different magnified single rod examples marked with a dashed line. (D, E) Additional single rod SIM z-projection images of WT IS-enriched retina sections immunolabeled with either (D) Rho-C-1D4 (magenta) or (E) Rho-N-4D2 (magenta); both co-immunolabeled with STX3 (cyan) and centrin-2 (yellow) antibodies. White arrows indicate Rho fluorescence that is colocalized with STX3 at the plasma membrane.

**Figure S3.** Alternate single rod STORM reconstruction examples from (A) Rho-GFP staining conditions co-labeled with STX3 (cyan) and centrin-2 yellow. Each example features plots of STORM molecule coordinates within the STX3+ IS hull for each channel and an adjacent plot with randomly plotted molecules within the IS hull (orange), as well as frequency and CDF graphs for distance to hull measurements from the plotted STORM coordinates. Molecule counts: (Ai) Rho-GFP n=9,634, STX3 n=7,643, Random n=9,634; (Aii) Rho-GFP n=4,298, STX3 n=14,385, Random n=4,298; (Aiii) Rho-GFP n=4,709, STX3 n=10,385, Random n=4,709. (B) STORM reconstructions of IS-enriched Rho-GFP-1D4/+ retina sections immunolabeled with NbGFP-A647 (cyan) and STX3 (cyan) and centrin-2 (yellow) antibodies. OSs are indicated. The centrin-2+ widefield images are superimposed on STORM reconstruction images. In single rod examples, the IS region is indicated, and the IS hull is outlined in cyan in a duplicate image. For each example Rho-GFP-1D4 and STX3 STORM molecule coordinates within the IS hull are plotted (top example: Rho-GFP-1D4 molecules = 8,135, STX3 molecules = 24,826; bottom example: Rho-GFP-1D4 molecules = 3,326, STX3 molecules = 17,765). In the adjacent plot, a random distribution of coordinates within the IS hull matching the number of Rho-GFP-1D4 molecules (4,239) are plotted in orange. Nearest distance to hull measurements for Rho-GFP-1D4, STX3 and random molecules are plotted in a frequency and CDF graphs. Colors in the graphs match the molecule plots. (C-F) Alternate single rod STORM reconstruction examples from (C) Rho-C-1D4, (D) Rho-N-4D2, (E) SNAP25 and (F) PDC conditions all co-labeled with STX3 (cyan) and centrin-2 (yellow) antibodies. Molecule counts (Ci) Rho-C-1D4 n=1,255, STX3 n=60,036, Random n=1,255; (Cii) Rho-C-1D4 n=9,586, STX3 n=37,569, Random n=9,586; (Ciii) Rho-C-1D4 n=1,152, STX3 n=66,161, Random n=1,152; (Di) Rho-N-4D2 n=4,965; STX3 n=67,836; Random n=4,965; (Dii) Rho-N-4D2 n=4,975; STX3 n=13,387; Random n=4,975; (E) SNAP25 n=7,697; STX3 n=25,196; Random n=7,697 (F) PDC n=10,920; STX3 n=18,619, Random n=10,920. Black arrows = Rho STORM molecules located at the STX3+ IS hull in rod examples where the majority of Rho molecules are internal (Aii, Cii, Dii).

**Figure S4.** (A, B) Violin plot graphs of STORM distance to hull normalized frequency values within (A) 0.1 µm and (B) 0.3 µm. N values are the same as in Figure 6F. Comparisons were tested for statistical significance using the Mann-Whitney U test. (A) PDC vs Rho-GFP **P-value = 0.0011; PDC vs Rho-GFP-1D4 ***P-value < 0.00001; PDC vs 1D4 ***P=0.0003; PDC vs 4D2 *P= 0.0172. (B) PDC vs Rho-GFP ***P-value = 0.0003; PDC vs Rho-GFP-1D4 ***P-value < 0.00001; PDC vs 1D4 **P= 0.0033. (C,D) Immunogold localization of syntaxin 3 and rhodopsin in mouse rods. (C) Single rod inner segment electron micrograph examples from a WT mouse retinas immunolabeled with STX3 antibody and nanogold secondary antibody. In a threshold image showing only the STX3+ immunogold particles, the approximate location of the IS plasma membrane is outlined in cyan. The outline is also continuous with the CC membrane. (D) Electron micrograph examples of rod ISs from IS-enriched WT mouse retinas immunolabeled with Rho-C-1D4. The IS plasma membrane is outlined in cyan in the threshold image. Some non-punctate staining from the BB and CC axoneme is present in the threshold images. mito = mitochondria.

**Figure S5.** (A) Alternate STORM single rod examples from Rho-GFP/+ IS-enriched retinas immunolabeled with NbGFP-A647 (magenta), and centrin and STX3 antibodies (combined, yellow), and Rab11a. (B) STORM examples for Rho-GFP-1D4/+ IS-enriched retinas immunolabeled with NbGFP-A647 (magenta), and centrin and STX3 antibodies (combined, yellow), and Rab11a. Rab11a+ clusters identified with Voronoi tessellation are in white. (C-D) STORM examples for Rho-GFP-1D4/+ IS-enriched retinas immunolabeled with NbGFP-A647 (magenta), and centrin and STX3 antibodies (combined, yellow), and either (C) DYNC1H1 or (D) Rootletin antibody.

## Notes

### Competing Interest Statement

The authors have declared no competing interest.

