## Supplemental Figures for "Mapping rhodopsin trafficking in rod photoreceptors with quantitative super-resolution microscopy"

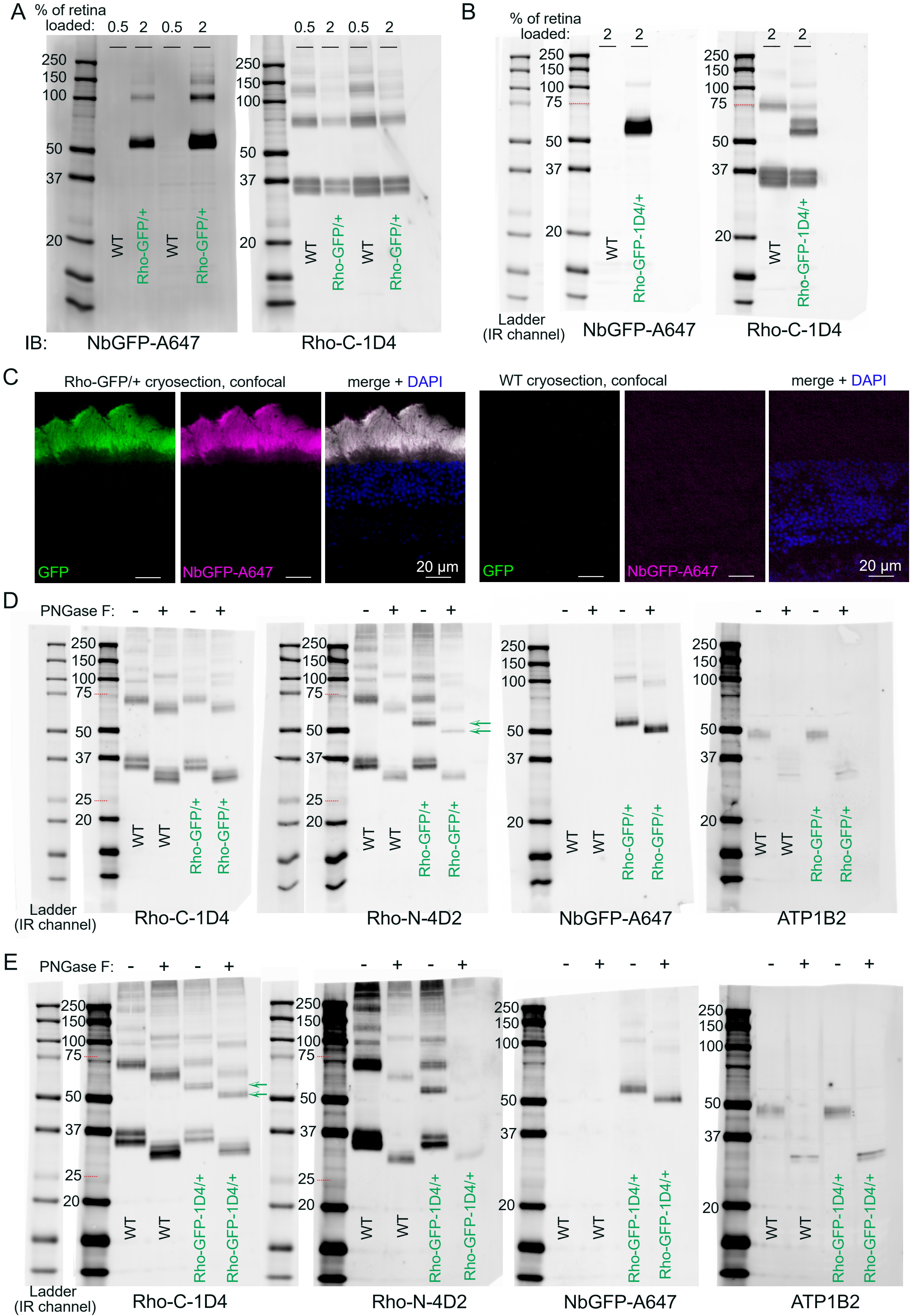
A
% of retina loaded:
0.5 2 0.5 2
250 150 100 50 37 20
WT Rho-GFP/+ WT Rho-GFP/+
IB:
NbGFP-A647
Rho-C-1D4
B
% of retina loaded:
2 2
250 150 100 75 50 37 20
WT Rho-GFP-1D4/+
Ladder (IR channel)
NbGFP-A647
Rho-C-1D4
C
Rho-GFP/+ cryosection, confocal
merge + DAPI
GFP
NbGFP-A647
20 µm
WT cryosection, confocal
merge + DAPI
GFP
NbGFP-A647
20 µm
D
PNGase F: - + - +
250 150 100 75 50 37 25 20
WT WT Rho-GFP/+ Rho-GFP/+
Ladder (IR channel)
Rho-C-1D4
Rho-N-4D2
NbGFP-A647
ATP1B2
E
PNGase F: - + - +
250 150 100 75 50 37 25 20
WT WT Rho-GFP-1D4/+ Rho-GFP-1D4/+
Ladder (IR channel)
Rho-C-1D4
Rho-N-4D2
NbGFP-A647
ATP1B2

**Figure S1.** (A, B) Western blot test of NbGFP-A647 immunolabeling specificity. (A) WT and Rho-GFP/+ retinal lysates used as either 0.5% or 2% total volume of 1 mouse retina. In Rho-GFP/+ lysates, a prominent and specific NbGFP+ band was found ~60 kDa corresponding to monomeric Rho-GFP protein. Both WT and Rho-GFP/+ lysates contained Rho-C-1D4-positive bands corresponding to endogenous mouse Rho protein. Mouse Rho protein levels are apparently lower in Rho-GFP/+ lysates – corresponding to the one WT Rho allele in these mice. In addition, Rho-C-1D4 antibody does not immunolabel Rho-GFP protein from Rho-GFP/+ lysates. IB=immunoblot condition. (B) WT and Rho-GFP-1D4/+ retinal lysates used as 2% total volume of 1 mouse retina. In Rho-GFP-1D4/+ lysates specific NbGFP-A647+ and Rho-C-1D4+ bands are found ~60 kDa corresponding to monomeric Rho-GFP-1D4 protein. As is (A), endogenous mouse Rho protein levels are lower in Rho-GFP-1D4/+ lysates. For all western blots, molecular weight marker sizes are indicated in kDa. (C) To test NbGFP-A647 immunolabeling specificity with immunofluorescence, 10 µm eye cup cryosections from Rho-GFP/+ mice (age P71) and from WT mice (age P60) were immunolabeled with NbGFP-A647 and counterstained with DAPI. NbGFP-647 specifically labels the outer segments (OS) in the Rho-GFP/+ section (magenta) and overlaps with GFP (green). No detectable NbGFP-A647 fluorescence was detected in the WT sections. Scalebar values match adjacent panels when not given. (D) Rho-GFP/+ or (E) Rho-GFP-1D4/+ adult retinal lysates treated with either Protein Deglycosylation Mix II from NEB (containing PNGase F) or buffer only. After treatment, 5 µg of total protein was loaded for SDS-PAGE, transfer and primary antibody labeling. A shift to lower molecular weight was observed for all Rho bands, including Rho-GFP bands from Rho-GFP/+ treated lysates probed with NbGFP-A647. In (D) Rho-GFP was also detected with Rho-N-4D2 immunolabeling; monomeric Rho-GFP bands are indicated with green arrows on this blot. In (E) Rho-GFP-1D4 was detected with Rho-C-1D4 immunolabeling and monomeric Rho-GFP-1D4 bands are indicated with green arrows.

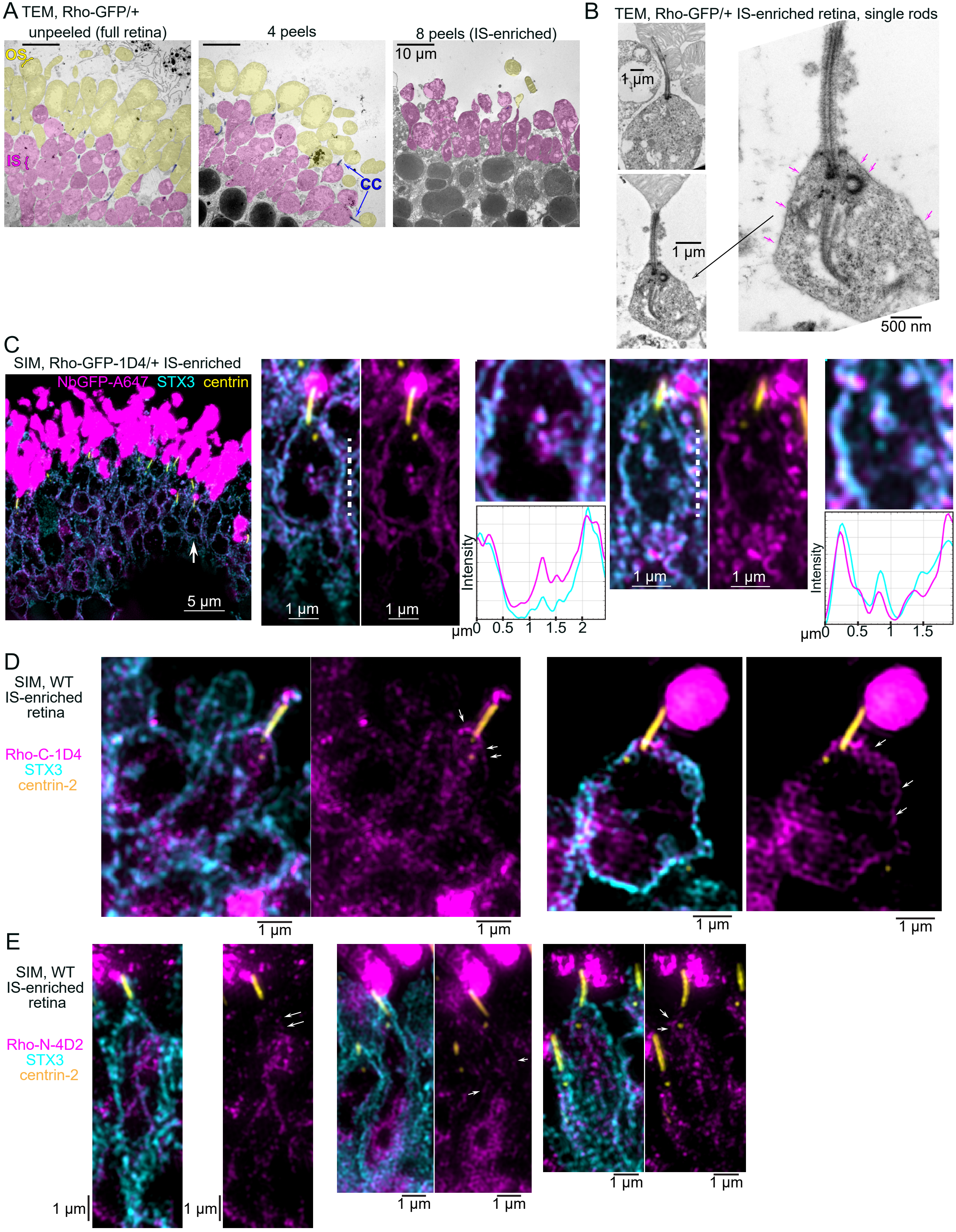

A TEM, Rho-GFP/+
unpeeled (full retina)
4 peels
8 peels (IS-enriched)
10 μm
OS
IS
CC
B TEM, Rho-GFP/+ IS-enriched retina, single rods
1 μm
1 μm
500 nm
C
SIM, Rho-GFP-1D4/+ IS-enriched
NbGFP-A647 STX3 centrin
5 μm
1 μm
1 μm
Intensity
μm 0 0.5 1 1.5 2
1 μm
1 μm
Intensity
μm 0 0.5 1 1.5
D
SIM, WT IS-enriched retina
Rho-C-1D4
STX3
centrin-2
1 μm
1 μm
1 μm
1 μm
E
SIM, WT IS-enriched retina
Rho-N-4D2
STX3
centrin-2
1 μm
1 μm
1 μm
1 μm
1 μm
1 μm

**Figure S2.** (A) TEM images of Rho-GFP/+ retina slices that were either unpeeled, peeled 4 times or peeled 8 times (the IS-enriched condition). Images were pseudocolored to point out key rod structures as follows: OS = yellow, IS = magenta, connecting cilia (CC)/basal bodies = blue. Scalebar values match adjacent panels when not labeled. (B) Alternate TEM single rod examples from Rho-GFP/+ IS-enriched retinas. The IS plasma membrane is annotated with magenta arrows. (C) SIM images from IS-enriched Rho-GFP/+ retinas immunolabeled for NbGFP-A647 (magenta), STX3 (cyan), and centrin (yellow). To demonstrate Rho-GFP colocalization with STX3 at the IS plasma membrane, row average intensity plots are shown for portions of the IS from 2 different magnified single rod examples marked with a dashed line. (D, E) Additional single rod SIM z-projection images of WT IS-enriched retina sections immunolabeled with either (D) Rho-C-1D4 (magenta) or (E) Rho-N-4D2 (magenta); both co-immunolabeled with STX3 (cyan) and centrin-2 (yellow) antibodies. White arrows indicate Rho fluorescence that is colocalized with STX3 at the plasma membrane.

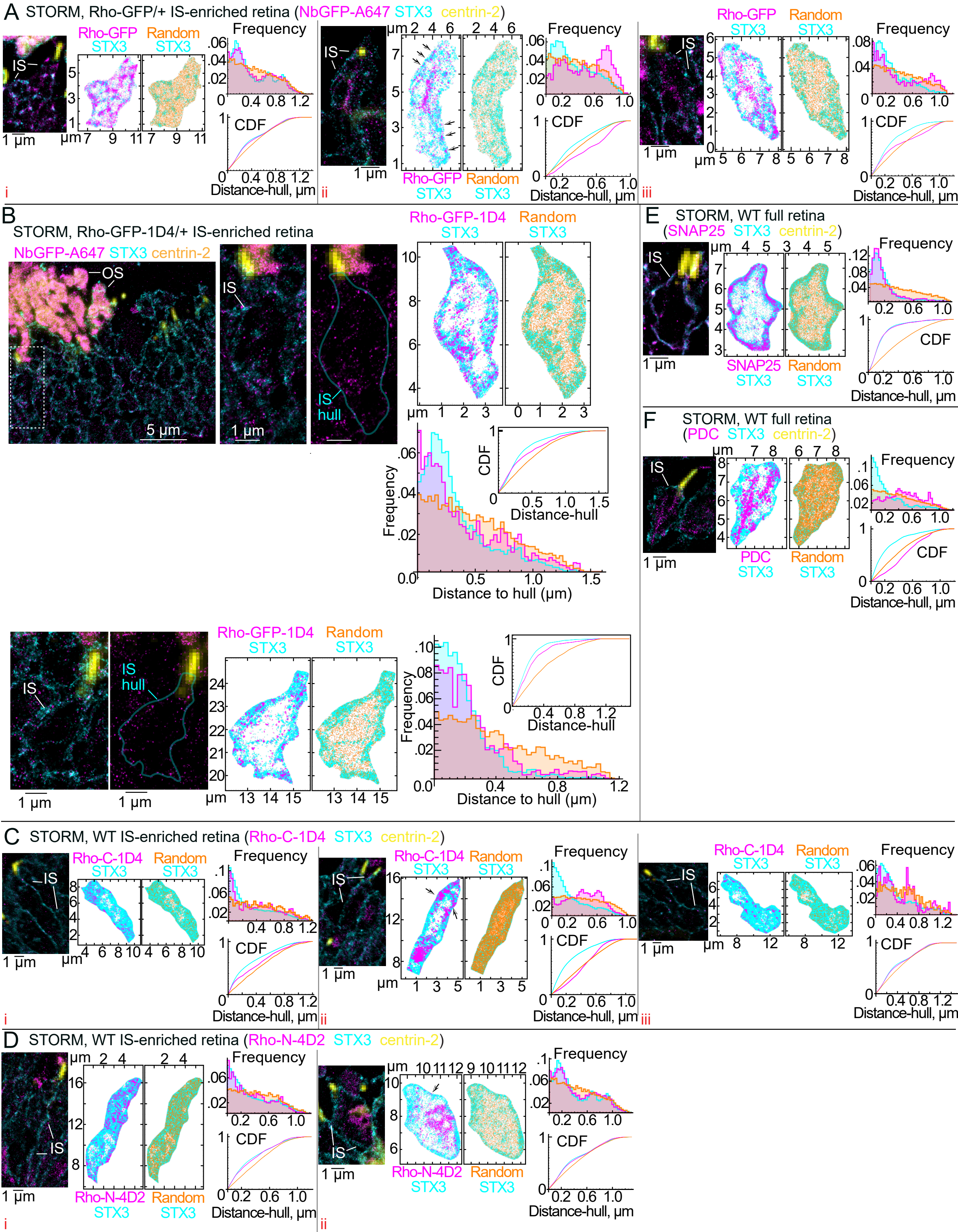

A STORM, Rho-GFP/+ IS-enriched retina (NbGFP-A647 STX3 centrin-2)
IS
Rho-GFP STX3
Random STX3
Frequency
CDF
Distance-hull, µm
1 µm
i
ii
iii
B
STORM, Rho-GFP-1D4/+ IS-enriched retina
NbGFP-A647 STX3 centrin-2
OS
5 µm
IS hull
Rho-GFP-1D4 STX3
Random STX3
Frequency
Distance to hull (µm)
Distance-hull
E STORM, WT full retina (SNAP25 STX3 centrin-2)
SNAP25 STX3
Random STX3
F STORM, WT full retina (PDC STX3 centrin-2)
PDC STX3
C STORM, WT IS-enriched retina (Rho-C-1D4 STX3 centrin-2)
Rho-C-1D4 STX3
D STORM, WT IS-enriched retina (Rho-N-4D2 STX3 centrin-2)
Rho-N-4D2 STX3

**Figure S3.** Alternate single rod STORM reconstruction examples from (A) Rho-GFP staining conditions co-labeled with STX3 (cyan) and centrin-2 yellow. Each example features plots of STORM molecule coordinates within the STX3+ IS hull for each channel and an adjacent plot with randomly plotted molecules within the IS hull (orange), as well as frequency and CDF graphs for distance to hull measurements from the plotted STORM coordinates. Molecule counts: (Ai) Rho-GFP n=9,634, STX3 n=7,643, Random n=9,634; (Aii) Rho-GFP n=4,298, STX3 n=14,385, Random n=4,298; (Aiii) Rho-GFP n=4,709, STX3 n=10,385, Random n=4,709. (B) STORM reconstructions of IS-enriched Rho-GFP-1D4/+ retina sections immunolabeled with NbGFP-A647 (cyan) and STX3 (cyan) and centrin-2 (yellow) antibodies. OSs are indicated. The centrin-2+ widefield images are superimposed on STORM reconstruction images. In single rod examples, the IS region is indicated, and the IS hull is outlined in cyan in a duplicate image. For each example Rho-GFP-1D4 and STX3 STORM molecule coordinates within the IS hull are plotted (top example: Rho-GFP-1D4 molecules = 8,135, STX3 molecules = 24,826; bottom example: Rho-GFP-1D4 molecules = 3,326, STX3 molecules = 17,765). In the adjacent plot, a random distribution of coordinates within the IS hull matching the number of Rho-GFP-1D4 molecules (4,239) are plotted in orange. Nearest distance to hull measurements for Rho-GFP-1D4, STX3 and random molecules are plotted in a frequency and CDF graphs. Colors in the graphs match the molecule plots. (C-F) Alternate single rod STORM reconstruction examples from (C) Rho-C-1D4, (D) Rho-N-4D2, (E) SNAP25 and (F) PDC conditions all co-labeled with STX3 (cyan) and centrin-2 (yellow) antibodies. Molecule counts (Ci) Rho-C-1D4 n=1,255, STX3 n=60,036, Random n=1,255; (Cii) Rho-C-1D4 n=9,586, STX3 n=37,569, Random n=9,586; (Ciii) Rho-C-1D4 n=1,152, STX3 n=66,161, Random n=1,152; (Di) Rho-N-4D2 n=4,965; STX3 n=67,836; Random n=4,965; (Dii) Rho-N-4D2 n=4,975; STX3 n=13,387; Random n=4,975; (E) SNAP25 n=7,697; STX3 n=25,196; Random n=7,697 (F) PDC n=10,920; STX3 n=18,619, Random n=10,920. Black arrows = Rho STORM molecules located at the STX3+ IS hull in rod examples where the majority of Rho molecules are internal (Aii, Cii, Dii).

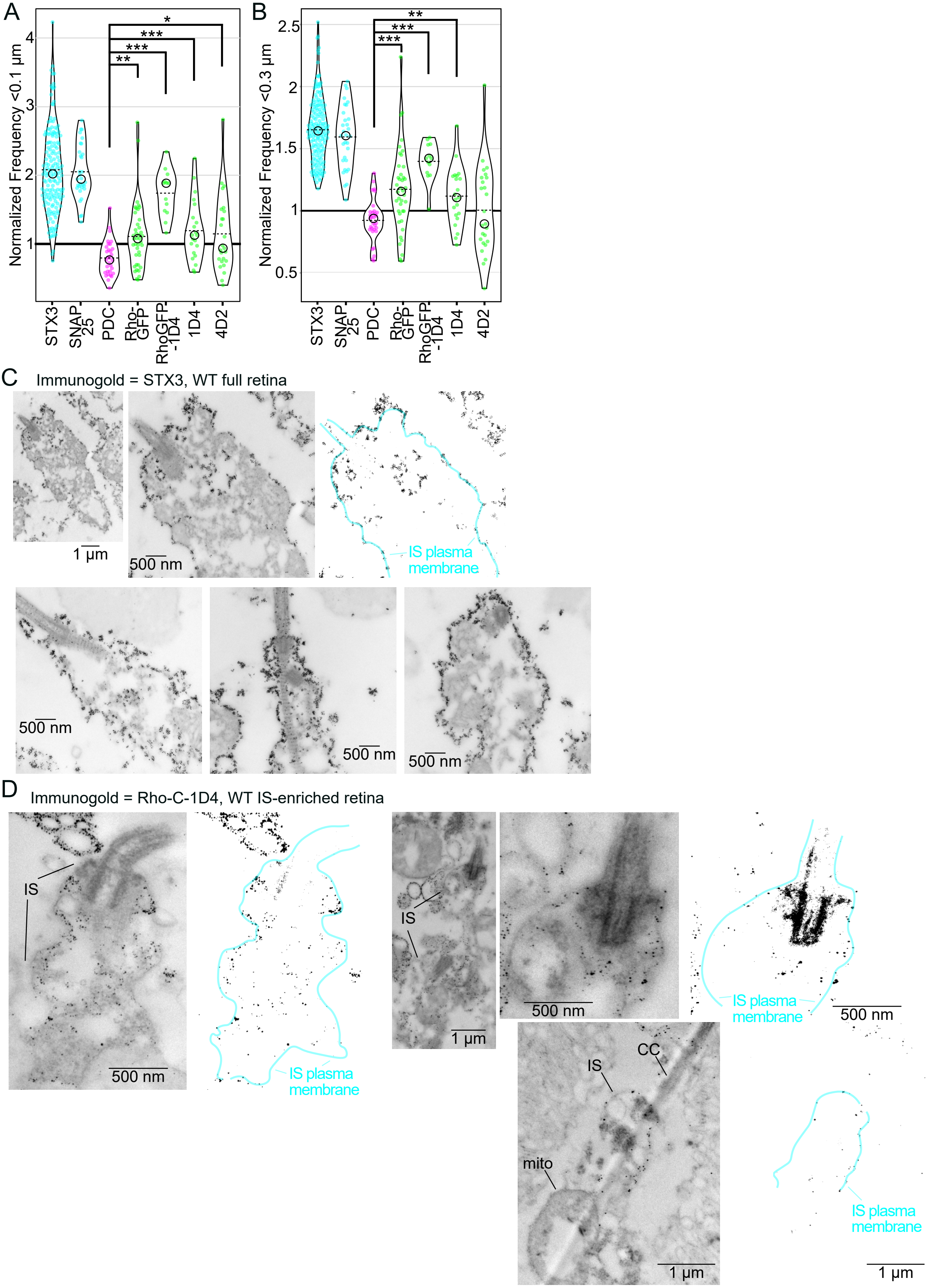
A
Normalized Frequency <0.1 µm
STX3
SNAP 25
PDC
Rho-GFP
RhoGFP -1D4
1D4
4D2
*
***
***
**
B
Normalized Frequency <0.3 µm
**
***
***
C
Immunogold = STX3, WT full retina
1 µm
500 nm
IS plasma membrane
D
Immunogold = Rho-C-1D4, WT IS-enriched retina
IS
CC
mito
IS plasma membrane
500 nm
1 µm

**Figure S4.** (A, B) Violin plot graphs of STORM distance to hull normalized frequency values within (A) 0.1 µm and (B) 0.3 µm. N values are the same as in Figure 6F. Comparisons were tested for statistical significance using the Mann-Whitney U test. (A) PDC vs Rho-GFP **P-value = 0.0011; PDC vs Rho-GFP-1D4 ***P-value < 0.00001; PDC vs 1D4 ***P=0.0003; PDC vs 4D2 *P= 0.0172. (B) PDC vs Rho-GFP ***P-value = 0.0003; PDC vs Rho-GFP-1D4 ***P-value < 0.00001; PDC vs 1D4 **P= 0.0033. (C,D) Immunogold localization of syntaxin 3 and rhodopsin in mouse rods. (C) Single rod inner segment electron micrograph examples from a WT mouse retinas immunolabeled with STX3 antibody and nanogold secondary antibody. In a threshold image showing only the STX3+ immunogold particles, the approximate location of the IS plasma membrane is outlined in cyan. The outline is also continuous with the CC membrane. (D) Electron micrograph examples of rod ISs from IS-enriched WT mouse retinas immunolabeled with Rho-C-1D4. The IS plasma membrane is outlined in cyan in the threshold image. Some non-punctate staining from the BB and CC axoneme is present in the threshold images. mito = mitochondria.

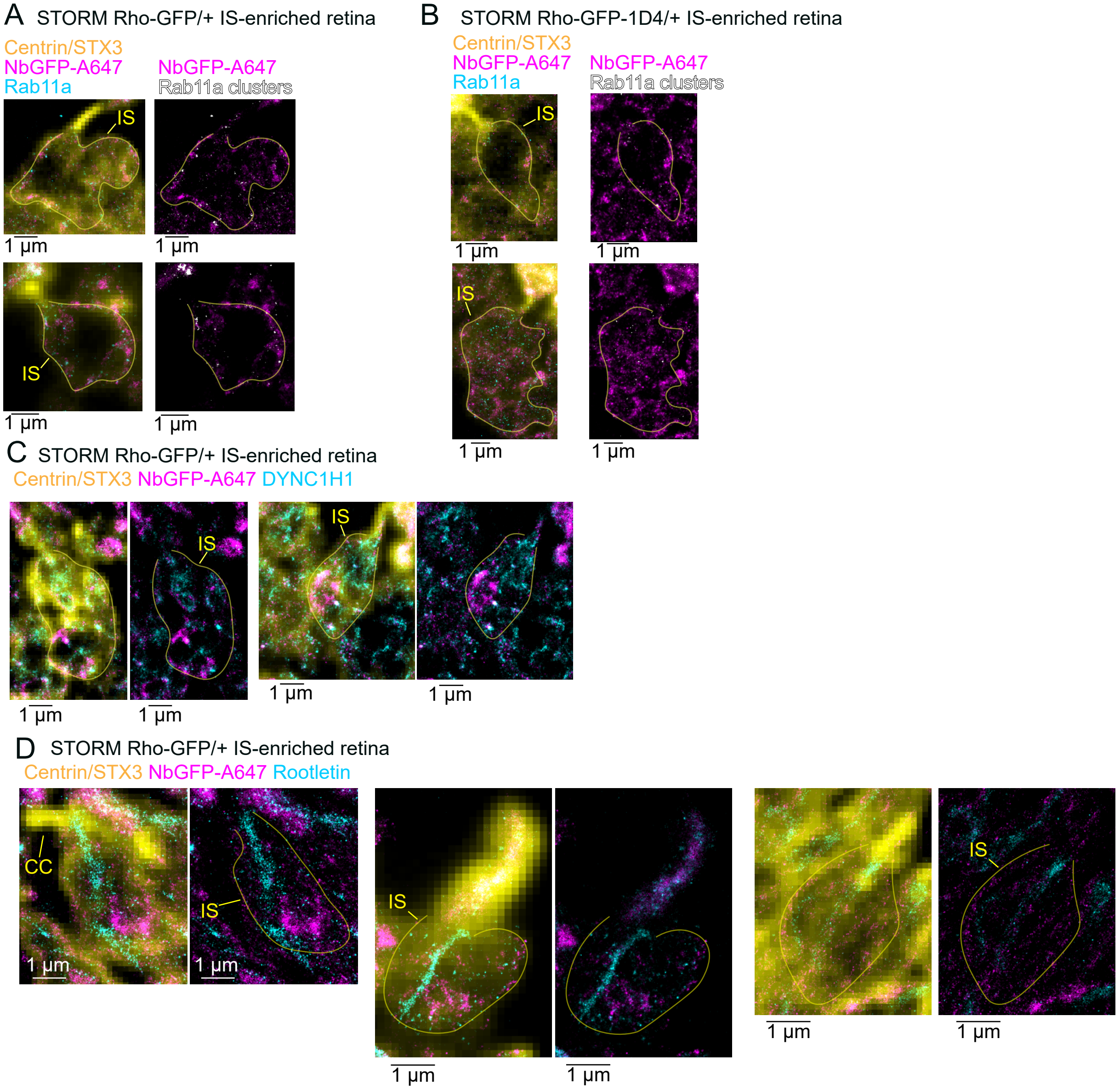


**Figure S5.** (A) Alternate STORM single rod examples from Rho-GFP/+ IS-enriched retinas immunolabeled with NbGFP-A647 (magenta), and centrin and STX3 antibodies (combined, yellow), and Rab11a. (B) STORM examples for Rho-GFP-1D4/+ IS-enriched retinas immunolabeled with NbGFP-A647 (magenta), and centrin and STX3 antibodies (combined, yellow), and Rab11a. Rab11a+ clusters identified with Voronoi tessellation are in white. (C-D) STORM examples for Rho-GFP-1D4/+ IS-enriched retinas immunolabeled with NbGFP-A647 (magenta), and centrin and STX3 antibodies (combined, yellow), and either (C) DYNC1H1 or (D) Rootletin antibody.
